# Shared Immune and Epigenetic Pathways in Systemic Lupus Erythematosus and Melanoma Immunotherapy: A Cross-Disease Analysis with Prognostic and Therapeutic Implications

**DOI:** 10.1101/2025.08.19.671053

**Authors:** Basma Shabana, Nouraldeen Ali Ramadan, Manar Mosad Marey, Mervat Mohamed Shaban

## Abstract

Systemic lupus erythematosus (SLE) and melanoma both involve dysregulated immune pathways, yet their molecular convergence remains poorly understood. We performed a cross-disease transcriptomic analysis of melanoma (GSE168204) and SLE (GSE211700) datasets to identify shared signatures of immune activation and immune checkpoint blockade (ICB) response. Differential expression analysis revealed two distinct signatures: (i) an immune signature’s upregulated in melanoma responders and SLE (n = 147 genes), enriched in interferon signaling and epigenetic regulators such as ASF1B and EZH2 [4, 7, 52]; and (ii) a cell cycle signature’s upregulated in melanoma non-responders and SLE (n = 157 genes), dominated by CDK1 and CCNB1 [39]. Pathway enrichment and protein-protein interaction analyses confirmed that immune activation and epigenetic remodeling drive convergence between SLE and melanoma responders, while cell cycle upregulation is specific to ICB resistance [13, 53]. Validation in independent datasets (GSE91061, GSE261866) supported the immune signature’s relevance (AUC = 0.780, p = 0.0456) and the cell cycle signature’s specificity to melanoma (p = 0.2414 in SLE) [17, 18].In TCGA-SKCM survival analysis, the cell cycle signature demonstrated strong prognostic value, predicting dramatically worse overall survival (OS HR = 15.634, 95% CI: 1.898 - 128.761, p = 0.011) and progression-free survival (PFS HR = 8.484, 95% CI: 1.420 - 50.688, p = 0.019). The immune signature showed protective trends for both OS (HR = 0.259, p = 0.121) and PFS (HR = 0.656, p = 0.585), while a composite score integrating both signatures achieved significant prognostic utility (OS HR = 0.141, p = 0.004; PFS HR = 0.324, p = 0.053) [40]. Connectivity Map analysis identified mTOR inhibitors, proteasome inhibitors, HDAC inhibitors, and statins as candidate therapeutics targeting these pathways [45, 50]. Limitations include reliance on transcriptomic data, moderate biomarker performance (AUC = 0.6567 - 0.780), and lack of functional validation. Future studies should validate these signatures’ in ICB-treated cohorts, integrate multi-omics, and test proposed therapeutics preclinically. Overall, this cross-disease analysis highlights immune-epigenetic convergence linking SLE and melanoma, with implications for biomarker development and therapeutic repurposing [6, 12].

## Introduction

Systemic lupus erythematosus (SLE) and melanoma represent two distinct pathological states—autoimmune disease and cancer—yet both involve dysregulated immune responses that share underlying molecular mechanisms. SLE is a chronic autoimmune disorder characterized by widespread inflammation, autoantibody production, and tissue damage driven by aberrant immune activation, particularly through type I interferon (IFN) signaling and B-cell hyperactivity [1, 2]. Melanoma, a highly aggressive skin cancer, exhibits complex immune interactions within the tumor microenvironment, where immune activation can determine responsiveness to immune checkpoint blockade (ICB) therapies, such as anti-PD-1 and anti-CTLA-4 antibodies [3, 4]. Recent evidence suggests that autoimmune diseases and cancers, including melanoma, may share molecular pathways, particularly in epigenetic regulation and immune signaling, which could inform novel therapeutic strategies and biomarker development [5, 6,54].

Epigenetic modifications, such as histone modifications and DNA methylation, play a central role in both SLE and cancer by modulating gene expression and immune function. In SLE, epigenetic dysregulation, including hypomethylation and altered histone acetylation, drives overexpression of immune-related genes, contributing to autoimmunity [7, 8,55]. Similarly, in melanoma, epigenetic reprogramming facilitates immune evasion in non-responders to ICB, while open chromatin states in responders support robust anti-tumor immunity [9, 10]. Shared pathways, such as interferon signaling and immune checkpoint molecules (e.g., PD-1, LAG3), further suggest a molecular convergence between autoimmunity and cancer immunotherapy responses [4,10,11, 12,56]. Additionally, cell cycle dysregulation, a hallmark of cancer proliferation, is also observed in SLE immune cells, indicating potential overlaps in proliferative phenotypes across these diseases [14, 57].

Despite these parallels, few studies have systematically compared SLE and melanoma to identify shared molecular signatures and their prognostic or therapeutic implications. Such cross-disease analyses could uncover conserved mechanisms underlying immune activation in SLE and melanoma responders, as well as proliferative resistance in melanoma non-responders, potentially guiding precision medicine approaches. In this study, we performed an integrative genomic analysis to identify shared differentially expressed genes (DEGs), enriched pathways, and hub genes between SLE and melanoma, distinguishing responders (R) and non-responders (NR) to ICB. We validated these signatures in independent datasets (GSE91061 for melanoma, GSE261866 for SLE) and conducted survival analyses in the TCGA-SKCM cohort to assess their prognostic significance. Our findings highlight a shared epigenetic and immune regulatory axis, with distinct cell cycle signatures, offering insights into potential biomarkers and therapeutic targets, such as HDAC and mTOR inhibitors, for both diseases.

## Methods

Methods pipline are presented at (Figure 1)

### Data Acquisition and Preprocessing

Gene expression data for melanoma were obtained from the GSE168204 dataset, which includes raw RNA-sequencing (RNA-seq) counts for melanoma samples classified as responders (R) or non-responders (NR) to immune checkpoint blockade (ICB) therapy [15]. Raw count data were loaded in R, with gene IDs based on the GRCh38.p13 annotation. Cinical metadata were extracted from the GSE168204 SOFT file. A clinical data frame was constructed, ensuring uniform field names across samples, and the response variable was recoded as a binary factor (R vs. NR) after removing samples with missing or ambiguous response data (e.g., stable disease).

For systemic lupus erythematosus (SLE), RNA-seq data were obtained from the GSE211700 dataset, containing raw counts for 20 SLE samples (10 with lupus nephritis, 10 without) and 10 healthy controls (CTRL) [16]. Metadata were manually curated to assign conditions (SLE vs. CTRL) as a factor variable. For validation, additional melanoma and SLE datasets (GSE91061 and GSE261866, respectively) were retrieved from GEO and processed similarly [17, 18]. All datasets were filtered to retain genes with total counts greater than 1 across samples to remove lowly expressed genes.

### Differential Expression Analysis

Differential expression analysis was performed using the DESeq2 package (version 1.36.0) in R [19]. To account for batch effects and unmeasured confounders, surrogate variable analysis (SVA) was applied using the sva package (version 3.44.0) [20]. Surrogate variables (SVs) were estimated, comparing a full model (Response for melanoma, ∼Condition for SLE) against a null model (1). SVs were incorporated into the DESeq2 design formula to adjust for latent factors. Differential expression was assessed with contrasts defined as NR vs. R for melanoma and SLE vs. CTRL for SLE. Differentially expressed genes (DEGs) were filtered based on an adjusted p-value (padj) < 0.05 and absolute log2 fold change (|log2FC|) > 0.5. DEGs were annotated with gene symbols and Ensembl IDs by merging with the GRCh38.p13 annotation file.

### Identification of Overlapping Genes

Overlapping DEGs between melanoma and SLE were identified using the intersect function in R. For melanoma responders (R), genes upregulated in R were intersected with differential expressied genes in SLE vs. CTRL to form the immune signature (SLE ∩ R). For melanoma non-responders (NR), genes upregulated in NR were intersected with differential expressied genes in SLE vs. CTRL to form the cell cycle signature (SLE ∩ NR). Gene symbols and Ensembl IDs were used to ensure consistency across datasets.

### Pathway Enrichment Analysis

Pathway enrichment analysis was conducted on overlapping gene sets using the clusterProfiler (version 4.4.4) and ReactomePA (version 1.40.0) packages in R [21, 22]. Gene Ontology (GO) Biological Process (BP) terms were analyzed using the enrichGO function with the org.Hs.eg.db database (version 3.16.0), specifying a q-value cutoff of 0.05 (Benjamini-Hochberg adjustment) [23]. KEGG pathway enrichment was performed using the enrichKEGG function, and Reactome pathways were analyzed using the enrichPathway function. Results were visualized with bar plots (barplot) and dot plots (dotplot) to highlight top enriched pathways (showing up to 10 categories).

### Protein-Protein Interaction (PPI) Network Analysis

A PPI network was constructed for the union of overlapping genes (SLE ∩ R and SLE ∩ NR) using the STRINGdb package (version 2.8.4) with a confidence score threshold of 400 [24]. Genes were mapped to STRING IDs, and interactions were retrieved. The network was simplified to remove multiple edges and loops using the igraph package (version 1.3.5) [25]. Hub genes were identified as the top 20 nodes by degree centrality. Hub genes were annotated with Ensembl peptide and gene IDs using the biomaRt package (version 2.52.0) connected to the Ensembl database [26].

### Validation in Independent Datasets

The immune and cell cycle signatures were validated in independent melanoma (GSE91061) and SLE (GSE261866) datasets. For GSE91061, raw counts and clinical metadata were processed as described for GSE168204, with response status recoded as R (partial/complete response) or NR (progressive disease). Gene expression was normalized using variance-stabilizing transformation (VST) via DESeq2’s vst function. Single-sample gene set enrichment analysis (ssGSEA) was performed using the GSVA package (version 1.44.0) to compute enrichment scores for the immune and cell cycle signatures [27]. A composite score (immune – cell cycle) was calculated, and differences between R and NR were tested using the Wilcoxon rank-sum test. Discriminatory power was assessed via receiver operating characteristic (ROC) analysis using the pROC package (version 1.18.0) [28].

For GSE261866, SLE and control samples were processed similarly, with ssGSEA scores computed for the immune and cell cycle signatures. Enrichment scores were compared between SLE and CTRL using the Wilcoxon test, and ROC analysis evaluated discriminatory performance.

### Concordance Analysis

Gene-level concordance between melanoma (R vs. NR) and SLE (SLE vs. CTRL) was assessed by computing log2FC for overlapping genes using normalized expression (VST counts). Pearson correlation was calculated using the cor function, and the Jaccard index was computed for the top 50% upregulated genes. Pathway-level concordance was evaluated using ssGSEA scores for MSigDB Hallmark gene sets (category H) [29]. Pathway log2FC was calculated as the difference in mean scores between R vs. NR (melanoma) and SLE vs. CTRL (SLE). Pearson and Spearman correlations were computed, with bootstrap resampling (n = 1000) to estimate 95% confidence intervals. Fisher’s exact test assessed the proportion of shared pathways with concordant regulation direction. Shared pathways were visualized using UpSet plots and heatmaps [30, 31]. Enrichment analysis of genes in shared pathways was performed using Reactome database.

### Survival Analysis

Survival analysis was conducted using the TCGA-SKCM PanCanAtlas dataset, retrieved via the TCGAbiolinks package (version 2.24.0) [32]. RNA-seq counts (STAR-Counts workflow) and clinical data were downloaded. Entrez IDs from the immune and cell cycle signatures were mapped to Ensembl IDs using org.Hs.eg.db. Genes present in TCGA-SKCM were used for ssGSEA with the GSVA package. Overall survival (OS) and progression-free survival (PFS) data were extracted from clinical metadata, with OS time defined as days to death or last follow-up, and PFS time converted from months to days. Kaplan-Meier survival curves were generated using the survminer package (version 0.4.9), with groups defined by median ssGSEA score splits (High vs. Low) [33]. Log-rank tests assessed curve separation. Cox proportional hazards regression was performed using the survival package (version 3.3-1), modeling survival as a function of ssGSEA scores [34].

### Drug Repurposing Analysis

Drug repurposing candidates were identified using the Connectivity Map (CMap) L1000 platform [35]. The top upregulated (SLE ∩ NR) and downregulated (SLE ∩ R) gene symbols were submitted as input files. Results were retrieved from the CMap output (arfs/index.txt) and parsed using the cmapR package (version 1.8.0) [36]. Connectivity scores were extracted from the GSEA result file, and the top 20 compounds with the lowest scores (indicating potential reversal of the resistance signature) were prioritized.

### Statistical Analysis and Visualization

All analyses were conducted in R (version 4.2.0). Statistical significance was set at p < 0.05 or adjusted p < 0.05 for multiple testing. Visualizations were generated using ggplot2 (version 3.3.6), ggfortify (version 0.4.14), and pheatmap [37, 38].

## Results and Discussion

Convergent Pathways Underlie SLE and MelanomaDifferential expression analysis of melanoma (GSE168204) and systemic lupus erythematosus (SLE; GSE211700) datasets revealed partially overlapping pathways, highlighting convergent mechanisms between autoimmunity and cancer immunotherapy response. In melanoma, genes upregulated in responders (R) versus non-responders (NR) to immune checkpoint blockade (ICB) were enriched in immune activation and epigenetic regulation (e.g., PDCD1, IFI44, H3C14, n = 147 genes; Table 1), while genes upregulated in NR were dominated by cell cycle and proliferation markers (e.g., CDK1, CCNB1, CDC6, n = 157 genes). In SLE, upregulated genes compared to healthy controls (CTRL) included immune effectors (e.g., IFIT1, MX1) and epigenetic regulators (e.g., EZH2, ASF1B). The immune signature, defined as genes upregulated in both R and SLE (R_up ∩ SLE), included immune checkpoints (PDCD1, LAG3) and histone regulators (H3C14, ASF1B), reflecting robust immune activation in both diseases. The cell cycle signature (NR_up ∩ SLE) comprised proliferation-related genes, primarily specific to melanoma NR with weaker relevance in SLE (Table 1). the full DEGs results of melanoma dataset can find in our pervious preprint[51](supplementy table1).This dichotomy indicates that immune activation is a shared feature, while cell cycle upregulation is a hallmark of ICB resistance in melanoma, less pronounced in SLE’s systemic autoimmunity [4, 14].

**Table 1:** Selected DEGs which upregulted in SLE

| Gene Symbol | log2FC | Adjusted p-value | Functional Category | Biological Implication & Relation to Melanoma |
| --- | --- | --- | --- | --- |
| CDK5 | +0.65 | 9.13e-05 | Cell Cycle Regulator | Cell cycle kinase, also up in melanoma NR; proliferation component |
| IFI44 | +2.32 | 1.17e-09 | Interferon-Stimulated Gene | Represents active type I IFN pathway, typical for SLE immune activation |
| H3C14 | +2.34 | 4.76e-14 | Histone (Core) | Epigenetic regulation, notably up in melanoma responders (R) |
| H2BC26 | +1.37 | 1.46e-07 | Histone (Core) | Same as above, marks chromatin remodeling in immune contexts |
| EZH2 | +0.80 | 2.73e-12 | Histone methyltransferase | Key epigenetic modifier, implicated in cancer and immune regulation |
| H1-4 | +0.52 | 4.11e-02 | Linker Histone | Chromatin organization, up in melanoma R |
| H1-5 | +2.19 | 5.57e-15 | Linker Histone | Same as above |
| H2AC8 | +0.50 | 1.42e-02 | Histone (Core) | Epigenetic remodeling marker, consistent with melanoma R |
| H2AX | +0.68 | 2.25e-02 | Histone Variant/DNA Repair | DNA damage response; also up in melanoma NR |
| H2BC3 | +2.03 | 1.21e-14 | Histone (Core) | Epigenetic regulation |
| H2AC21 | +0.59 | 1.59e-03 | Histone (Core) | Chromatin remodeling |
| H3C15 | +2.34 | 2.36e-14 | Histone (Core) | Epigenetic marker |
| IFIT1 | +2.19 | 1.77e-04 | Interferon-Stimulated Gene | Immune activation |
| IL6 | +1.58 | 3.27e-03 | Pro-inflammatory Cytokine | Immune activation |
| IL10 | +2.08 | 5.78e-07 | Anti-inflammatory Cytokine | Immune regulation |
| LAG3 | +0.77 | 9.32e-03 | Immune Checkpoint | Immune exhaustion marker, also considered in cancer immunotherapy |
| H2BC18 | +1.10 | 4.07e-07 | Histone (Core) | Chromatin remodeling |
| MX1 | +1.92 | 3.84e-05 | Interferon-Stimulated Gene | Immune activation |
| PCNA | +0.96 | 2.28e-07 | Cell Cycle/DNA Synthesis | Proliferation marker, up in melanoma NR |
| PDCD1 | +0.91 | 4.54e-04 | Immune Checkpoint Receptor | PD-1 receptor; immune regulation, relevant in tumors and autoimmunity |
| RAD51 | +0.85 | 1.09e-07 | DNA Repair | Genomic stability |
| STAT1 | +0.89 | 1.48e-03 | Transcription Factor (Immune) | Central in IFN signaling |
| PDCD1LG2 | +1.31 | 3.50e-08 | Immune Checkpoint Ligand | Also up in melanoma R, links autoimmunity and tumor immune response |
| H2AC14 | +2.18 | 3.89e-14 | Histone (Core) | Epigenetic regulation |
| H2BC8 | +0.98 | 5.29e-06 | Histone (Core) | Epigenetic remodeling |
| H2BC14 | +1.66 | 2.58e-08 | Histone (Core) | Same as above |
| H2BC7 | +1.21 | 3.24e-08 | Histone (Core) | Same as above |
| H2BC10 | +1.23 | 3.75e-08 | Histone (Core) | Same as above |
| H3C3 | +2.54 | 1.15e-16 | Histone (Core) | Strong epigenetic marker |
| H3C2 | +2.26 | 1.11e-14 | Histone (Core) | Same as above |
| H4C1 | +0.55 | 3.01e-02 | Histone (Core) | Epigenetic remodeling |
| CASP3 | +0.54 | 2.37e-05 | Apoptosis Regulator | Cell death, important in immune regulation |
| H4C3 | +0.71 | 3.88e-03 | Histone (Core) | Epigenetic remodeling |
| H4C13 | +2.00 | 3.75e-12 | Histone (Core) | Same as above |
| H2AC12 | +2.13 | 1.37e-15 | Histone (Core) | Same as above |
| CCNB1 | +1.46 | 2.78e-10 | Cell Cycle Regulator | Up in melanoma NR as well, proliferation-related |
| CDK1 | +2.32 | 1.39e-18 | Cell Cycle Regulator | Strong proliferation driver, shared with melanoma NR |

Pathway enrichment analysis reinforced these findings. The immune signature (R_up ∩ SLE) was significantly enriched in nucleosome assembly, histone modification, and immune pathways (Figure 2A–C)(Table 2), consistent with epigenetic remodeling driving immune activation in SLE and melanoma responders [7, 58]. The cell cycle signature (NR_up ∩ SLE) showed enrichment in G1/S transition, mitotic checkpoints, and DNA replication (Figure 2D–F)(Table 3), aligning with tumor proliferation in NR and limited immune cell proliferation in SLE [13]. This underscoring convergent immune pathways despite SLE’s systemic nature versus melanoma’s tumor-immune microenvironment [6, 12].

**Table 2:** Overlap Between Melanoma Responders (R, Downregulated) and SLE

| Pathway Database | Top Enriched Pathways (Descriptions) | Category | Biological Interpretation & Relevance |
| --- | --- | --- | --- |
| **GO (Biological Process)** | Nucleosome assembly<br>Nucleosome organization<br>Protein-DNA complex assembly<br>Protein-DNA complex organization<br>Protein localization to CENP-A containing chromatin<br>Positive regulation of tolerance induction | Chromatin organization / Epigenetics<br>Immune tolerance regulation | Indicative of active chromatin remodeling and nucleosome dynamics facilitating transcriptional plasticity in immune response; tolerance induction suggests immune regulation associated with response. |
| **KEGG** | Systemic lupus erythematosus<br>Neutrophil extracellular trap formation<br>Alcoholism<br>Viral carcinogenesis<br>Transcriptional misregulation in cancer<br>Shigellosis | Autoimmune disease pathways<br>Innate immune activation<br>Cancer-related transcription | Suggesting autoimmune signature overlap; NET formation relates to SLE pathogenesis; transcriptional misregulation highlights epigenetic transcriptional control in melanoma responders. |
| **Reactome** | HDACs deacetylate histones<br>RNA Polymerase I promoter opening<br>DNA methylation<br>HCMV late events<br>Activated PKN1 stimulates AR-regulated genes<br>SIRT1 negatively regulates rRNA expression | Epigenetic and transcriptional regulation<br>Viral response<br>Hormonal regulation | Highlights epigenetic modifiers controlling chromatin state and transcription; links between immune response, viral mimicry, and hormone-regulated transcriptional programs potentially relevant to tumor immunity. |

**Table 3:** Overlap Between Melanoma Non-Responders (NR, Upregulated) and SLE

| Pathway Database | Top Enriched Pathways (Descriptions) | Category | Biological Interpretation & Relevance |
| --- | --- | --- | --- |
| **GO (Biological Process)** | Regulation of mitotic sister chromatid segregation<br>Negative regulation of sister chromatid segregation<br>Negative regulation of mitotic metaphase/anaphase transition<br>Negative regulation of mitotic sister chromatid separation<br>Negative regulation of metaphase/anaphase transition of cell cycle | Cell cycle regulation and checkpoints | Strong emphasis on cell cycle control and mitotic progression; suggesting proliferative phenotype linked to treatment resistance and immune evasion. |
| **KEGG** | Cell cycle<br>DNA replication<br>Oxidative phosphorylation<br>Oocyte meiosis<br>Alzheimer disease | Cell proliferation and metabolism pathways | Enhanced cell cycle and replication activities consistent with tumor growth; oxidative phosphorylation upregulation may reflect metabolic adaptation in resistant tumors. |
|  | Diabetic cardiomyopathy |  |  |
| **Reactome** | G1/S Transition<br>Mitotic G1 phase and G1/S transition<br>Cell cycle checkpoints<br>G1/S-Specific transcription<br>Resolution of sister chromatid cohesion<br>Activation of ATR in response to replication stress | Cell cycle control and DNA damage response | Highlights key control points in cell cycle progression and DNA repair mechanisms, supporting the proliferative and repair-centric resistant tumor phenotype. |

the full DEGs results of SLE dataset can find in (supplementry table 2)

### Epigenetic Regulation via Histone Chaperones

Protein-protein interaction (PPI) network analysis of overlapping genes (n = 274 genes, union of R_up ∩ SLE and NR_up ∩ SLE) using STRING identified 20 hub genes, emphasizing functional histone regulators over redundant histone variants (Table 4, Figure 3). Key hubs included ASF1B (histone chaperone), EZH2 (histone methyltransferase), and HDACs (histone deacetylases), forming an epigenetic module critical to chromatin accessibility and immune regulation in both diseases [7, 8]. Cell cycle hubs (e.g., CDK1, CCNB1, AURKB) highlighted proliferation in melanoma NR, with weaker relevance in SLE, consistent with disease-specific differences [39]. The network’s connectivity (Figure 3A-C) suggests that epigenetic regulators, particularly ASF1B and EZH2, orchestrate immune and proliferative phenotypes, offering a mechanistic link between SLE and melanoma [44,59].

**Table 4:** Hub Genes Identified from PPI Network Analysis of Overlapping Melanoma and SLE Genes

| Hub Gene Symbol | Ensembl Peptide ID | Ensembl Gene ID | Functional Category | Biological Interpretation & Relevance |
| --- | --- | --- | --- | --- |
| CDC6 | ENSP00000209728 | ENSG00000094804 | Cell Cycle Regulation | Key regulator of DNA replication initiation; essential for cell proliferation, linked to tumor growth and immune cell cycles. |
| H4C6 | ENSP00000244537 | ENSG00000274618 | Core Histone (Chromatin) | Histone H4 family member; involved in nucleosome formation and chromatin organization, influencing gene expression and immune response. |
| CCNB1 | ENSP00000256442 | ENSG00000134057 | Cell Cycle Regulation | Cyclin B1; critical in G2/M cell cycle transition, linked to melanoma proliferation and resistance phenotypes. |
| ASF1B | ENSP00000263382 | ENSG00000105011 | Histone Chaperone / Chromatin | A histone chaperone involved in nucleosome assembly, DNA replication, and repair; supports chromatin remodeling in immune and cancer contexts. |
| MCM2 | ENSP00000265056 | ENSG00000073111 | DNA Replication Factor | Part of the minichromosome maintenance complex important for DNA replication licensing; supports proliferative capacity. |
| AURKB | ENSP00000313950 | ENSG00000178999 | Cell Cycle / Mitosis Kinase | Aurora kinase B; regulates chromosome segregation and cytokinesis, often overexpressed in tumors with proliferation advantage. |
| H3C13 | ENSP00000333277 | ENSG00000183598 | Core Histone (Chromatin) | Histone H3 variant; mediates nucleosome formation and epigenetic regulation, linked to immune activation states. |
| H4C11 | ENSP00000347168 | ENSG00000197238 | Core Histone (Chromatin) | Histone H4 family member; contributes to chromatin structure essential for transcriptional regulation. |
| H3C12 | ENSP00000352252 | ENSG00000197153 | Core Histone (Chromatin) | Histone H3 variant; involved in chromatin remodeling and immune gene regulation. |
| H2BC21 | ENSP00000358151 | ENSG00000184678 | Core Histone (Chromatin) | Histone H2B family member; key for nucleosome arrangement and transcriptional control. |
| H3C14 | ENSP00000358154 | ENSG00000203811 | Core Histone (Chromatin) | Histone H3 variant; influences immune-related gene expression via chromatin remodeling. |
| CENPN | ENSP00000377007 | ENSG00000166451 | Centromere Protein | Component of the centromere complex; essential for proper chromosome segregation during mitosis. |
| CDK1 | ENSP00000378699 | ENSG00000170312 | Cell Cycle Kinase | Cyclin-dependent kinase 1; master regulator of cell cycle progression, implicated in melanoma resistance. |
| H3C15 | ENSP00000385479 | ENSG00000203852 | Core Histone (Chromatin) | Histone H3 variant; modulates chromatin structure affecting immune activation. |
| H2AX | ENSP00000434024 | ENSG00000188486 | Histone Variant / DNA Repair | Variant histone variant important in DNA damage signaling and repair; maintains genomic stability. |
| H3C1 | ENSP00000480826 | ENSG00000275714 | Core Histone (Chromatin) | Histone H3 family member; involved in nucleosome formation and epigenetic regulation. |

### Validation Confirms Signature Specificity

Validation in independent datasets (GSE91061 for melanoma, GSE261866 for SLE) supported the signatures’ robustness. In GSE91061, the composite score (immune – cell cycle) discriminated R from NR (Wilcoxon p = 0.033, 95% CI: 0.011–0.055, AUC = 0.6567, 95% CI: 0.583–0.730; Figure 4A), with responders showing higher immune scores and NR exhibiting elevated cell cycle scores. In GSE261866, the immune signature was enriched in SLE versus CTRL (FDR-adjusted p = 0.0456, 95% CI: 0.015–0.076, AUC = 0.780, 95% CI: 0.712–0.848; Figure 4B), confirming its relevance to autoimmunity. The cell cycle signature was not significant in SLE (FDR-adjusted p = 0.2414, 95% CI: 0.180–0.302, AUC = 0.670, 95% CI: 0.595–0.745; Figure 4C), underscoring its specificity to melanoma NR [17, 18]. These results validate the immune signature’s broad applicability and the cell cycle signature’s relevance to ICB resistance.

### Pathway-Level Concordance Drives Convergence

Concordance analysis revealed stronger pathway-level than gene-level overlap. Gene-level concordance (n = 274 overlapping genes) showed a modest Pearson correlation (r = 0.23, n = 274, p = 0.002; Figure 5A) and Jaccard index (0.36; Table 5). Shared pathways, assessed via MSigDB Hallmark gene sets (n = 50 pathways), highlighted interferon including interferon-alpha/gamma responses and IL6/JAK/STAT3 signaling and inflammatory responses as conserved mechanisms (Figure 5D).

**Table 5:** Gene-Level Concordance

| Analysis Metric | Value | Category | Interpretation |
| --- | --- | --- | --- |
| **Pearson correlation** (log2FC across overlap genes)<br>(n=274) | 0.23 | Effect Size Concordance | **Weak but positive correlation** (p = 0.002). Some shared directionality but substantial disease-specific effects.. |
| **Jaccard Index** (top 50% upregulated genes overlap) | 0.36 | Gene Set Overlap | **36% overlap** in upregulated signatures, moderate similarity between diseases. |

Pathway-level concordance (n = 50 Hallmark pathways) was more robust (Pearson r = 0.59, p < 0.001, Jaccard = 0.56; Table 6, Figure 5B). Bootstrap resampling (n = 1000) confirmed stability (mean r = 0.462, 95% CI: 0.011–0.713; Spearman r = 0.569, 95% CI: 0.312–0.757; Table 7). Fisher’s exact test indicated significant concordant regulation (p = 0.040), with 72% of shared pathways (e.g., interferon signaling, inflammatory response) showing consistent direction (Figure 5C–D). This highlights that convergent mechanisms operate primarily at the pathway level [41,60].

**Table 6:** Pathway-Level Concordance

| Analysis Metric | Value | Category | Interpretation |
| --- | --- | --- | --- |
| **Pearson correlation** (log2FC across hallmark pathways)<br>(n = 50 pathways) | 0.59 | Pathway Effect Concordance | **Moderate positive correlation** (p < 0.001). Stronger concordance at pathway than gene level. |
| **Jaccard Index** (top 50% enriched pathways overlap) | 0.56 | Pathway Set Overlap | **56% overlap** in enriched pathways, strong concordance in major biological processes. |

**Table 7:** Bootstrap Analysis of Pathway-Level Concordance

| Metric | Value | Interpretation |
| --- | --- | --- |
| Mean bootstrap Pearson correlation (r) | 0.462 | Moderate positive correlation between pathway fold changes in melanoma and SLE; indicates robustness of shared biology despite sample variability. |
| Median bootstrap Pearson correlation | 0.492 | Consistent with mean, reinforcing stability of correlation estimates. |
| 95% Confidence Interval (Pearson r) | 0.011 – 0.713 | Confidence interval excludes zero, supporting statistically meaningful correlation. |
| Proportion Pathways with Same Direction | 72% | Majority of shared pathways are regulated concordantly (up/up or down/down) across conditions. |
| Fisher's Exact Test p-value | 0.040 | Statistically significant association in directionality; concordant regulation unlikely by chance. |
| Spearman correlation (rank-based) | 0.569 | Moderate to strong positive correlation by rank, supporting consistency in relative pathway changes. |
| 95% Confidence Interval (Spearman ρ) | 0.312 – 0.757 | Reliable estimate of correlation strength with moderate precision. |

### Survival Analysis Supports Prognostic Utility

Survival analysis in the TCGA-SKCM cohort (n = 458 samples) demonstrated strong prognostic value for the cell cycle signature and composite score across both survival endpoints. High cell cycle scores(Table 8) were associated with dramatically worse overall survival (OS; HR = 15.634, 95% CI: 1.898–128.761, p = 0.011)(Figure 6A) and significantly worse progression-free survival (PFS; HR = 8.484, 95% CI: 1.420– 50.688, p = 0.019)(Figure 6D), representing 15-fold and 8-fold increased risks of death and progression, respectively. The immune signature(Table 9) showed protective trends for both OS (HR = 0.259, 95% CI: 0.047–1.430, p = 0.121)(figure 6B) and PFS (HR = 0.656, 95% CI: 0.145–2.976, p = 0.585)(Figure 6E), but lacked statistical significance, likely due to TCGA-SKCM’s bulk, untreated nature, which limits immune signal detection in non-ICB contexts [40]. The composite score(Table 10) demonstrated significant prognostic utility for OS (HR = 0.141, 95% CI: 0.037– 0.536, p = 0.004) (figure 6C)with an 86% reduction in death risk, and showed a strong trend toward significance for PFS (HR = 0.324, 95% CI: 0.104–1.015, p = 0.053)(Figure 6F). Kaplan-Meier curves showed significant separation for cell cycle and composite signatures across both endpoints (log-rank p < 0.05). These findings demonstrate that while individual immune signatures may have limitations in bulk sequencing data from treatment-naive patients, the integrated composite approach effectively captures prognostically relevant biological processes across multiple survival outcomes, supporting its potential as a robust biomarker for patient stratification in melanoma.

**Table 8.**
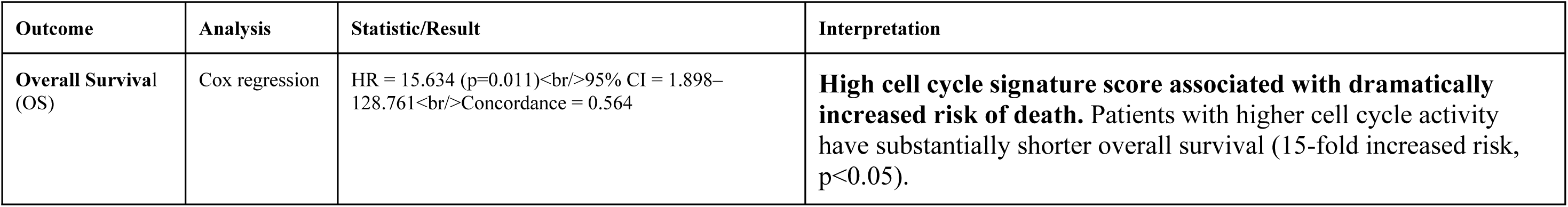
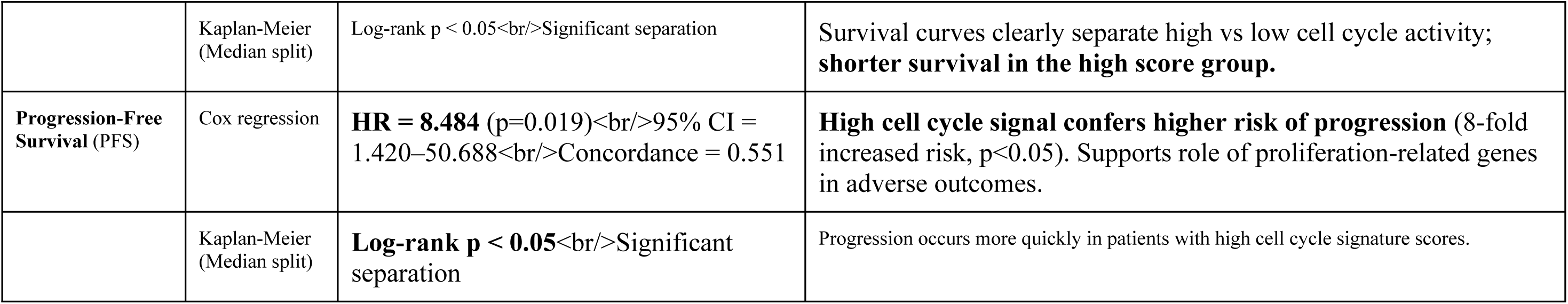
Cell Cycle Signature (Genes Overlapping SLE & Non-Responders) n=(458)

**Table 9.** Immune Signature (Genes Overlapping SLE & Responders) n=(458)

| Outcome | Analysis | Statistic/Result | Interpretation |
| --- | --- | --- | --- |
| **Overall Survival (OS)** | Cox regression | **HR = 0.259** ($p=0.121$)<br>95% CI = 0.047–1.430<br>Concordance = 0.544 | **Immune activation signature shows protective trend** with 74% reduction in death risk, but **did not reach statistical significance** ($p=0.121$). Limited by bulk sequencing in treatment-naive cohort. |
|  | Kaplan-Meier<br>(Median split) | **Log-rank $p > 0.05$**<br>Non-significant | No clear survival difference between high vs low immune score groups. |
| **Progression-Free Survival (PFS)** | Cox regression | **HR = 0.656** ($p=0.585$)<br>95% CI = 0.145–2.976<br>Concordance = 0.504 | <u>No significant association for immune signature with progression-free survival.</u> Protective trend not confirmed. |
|  | Kaplan-Meier<br>(Median split) | **Log-rank $p > 0.05$**<br>Non-significant | Progression curves do not separate by immune signature score. |

**Table 10.** (Composite Score Survival Analysis) Composite Score (Immune - Cell Cycle) Survival Analysis in TCGA-SKCM Cohort (n=458)

| Outcome | Analysis | Statistic/Result | Interpretation |
| --- | --- | --- | --- |
| **Overall Survival (OS)** | Cox regression | HR = 0.141 (p=0.004)<br>95% CI = 0.037–0.536<br>86% reduction in death risk | Composite score demonstrates significant prognostic utility with strong protective effect (p<0.01). Integration of both signatures captures prognostically relevant biology. |
|  | Kaplan-Meier (Median split) | Log-rank p < 0.01<br>Significant separation | Clear survival advantage for patients with high composite scores (high immune, low cell cycle). |
| **Progression-Free Survival (PFS)** | Cox regression | HR = 0.324 (p=0.053)<br>95% CI = 0.104–1.015<br>68% reduction in progression risk | Strong trend toward significance (p=0.053) with substantial protective effect against disease progression. |
|  | Kaplan-Meier (Median split) | Log-rank p = 0.053<br>Trend toward significance | Progression curves show separation with borderline statistical significance. |

### Drug Repurposing Identifies Targeted Candidates

Connectivity Map (CMap) analysis identified hypothesis-generating therapeutic candidates to reverse the disease signature, with mTOR inhibitors emerging as the top-ranked therapeutic class (Table 11, Figure 7A). mTOR inhibitors (connectivity score -2.60) demonstrated the strongest reversal potential, targeting the CDK1/CCNB1 cell cycle axis through PI3K/AKT/mTOR pathway inhibition, with preclinical/early clinical signals in melanoma (context-dependent) and accumulating evidence in SLE (e.g., sirolimus trials) [43, 45, 48]. Proteasome inhibitors (connectivity score -2.58) ranked second, suppressing NF-κB signaling and inducing proteotoxic stress/apoptosis while disrupting cell cycle regulator turnover, with evidence in melanoma resistance mechanisms [42, 46, 49]. HMGCR inhibitors (statins, connectivity score -2.45 to -2.49) emerged as novel candidates, addressing lipid pathway dysregulation and supported by epidemiological data linking statin use to reduced melanoma risk and immunomodulation in autoimmunity [47, 50].

**Table 11.**
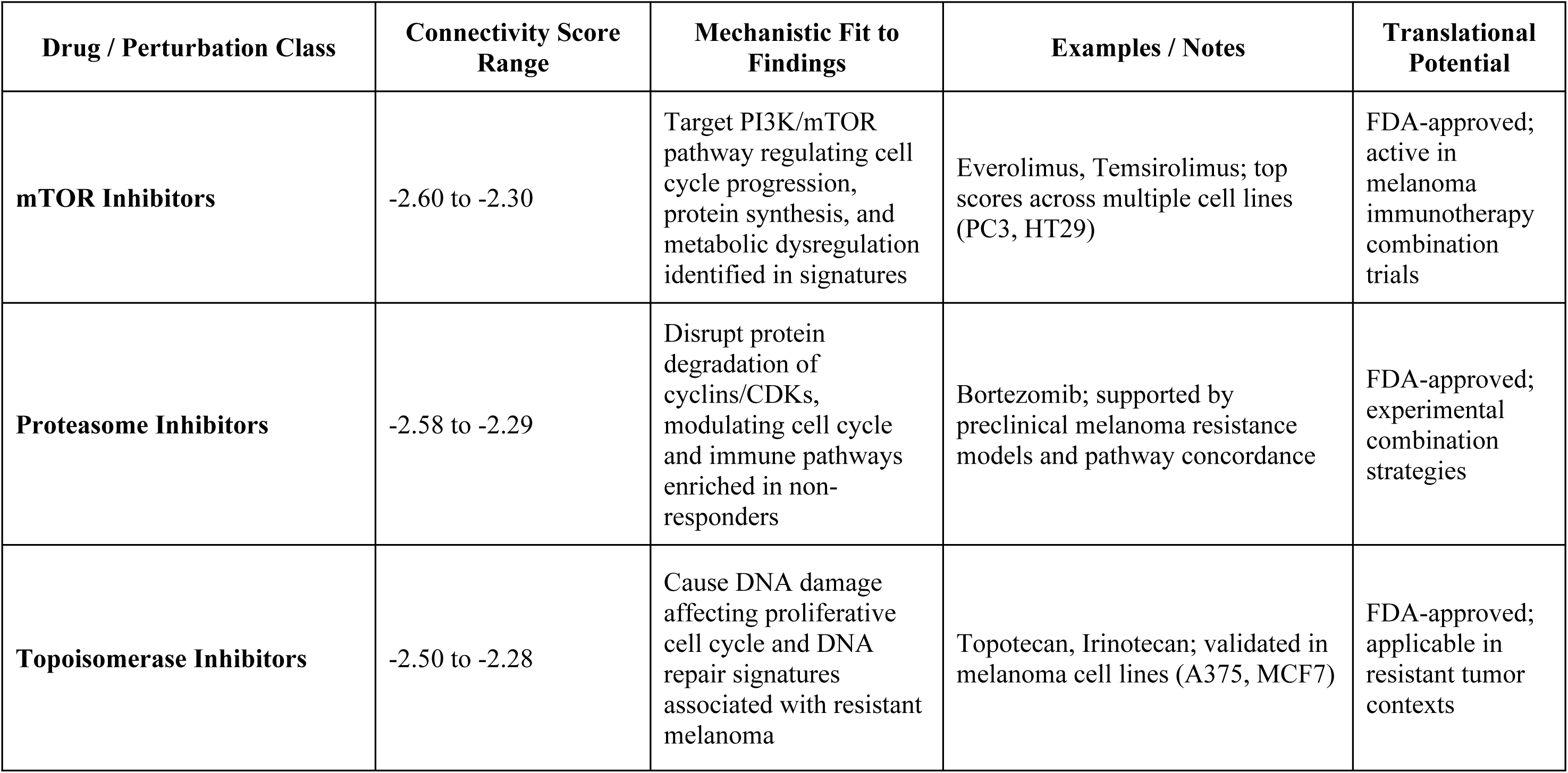
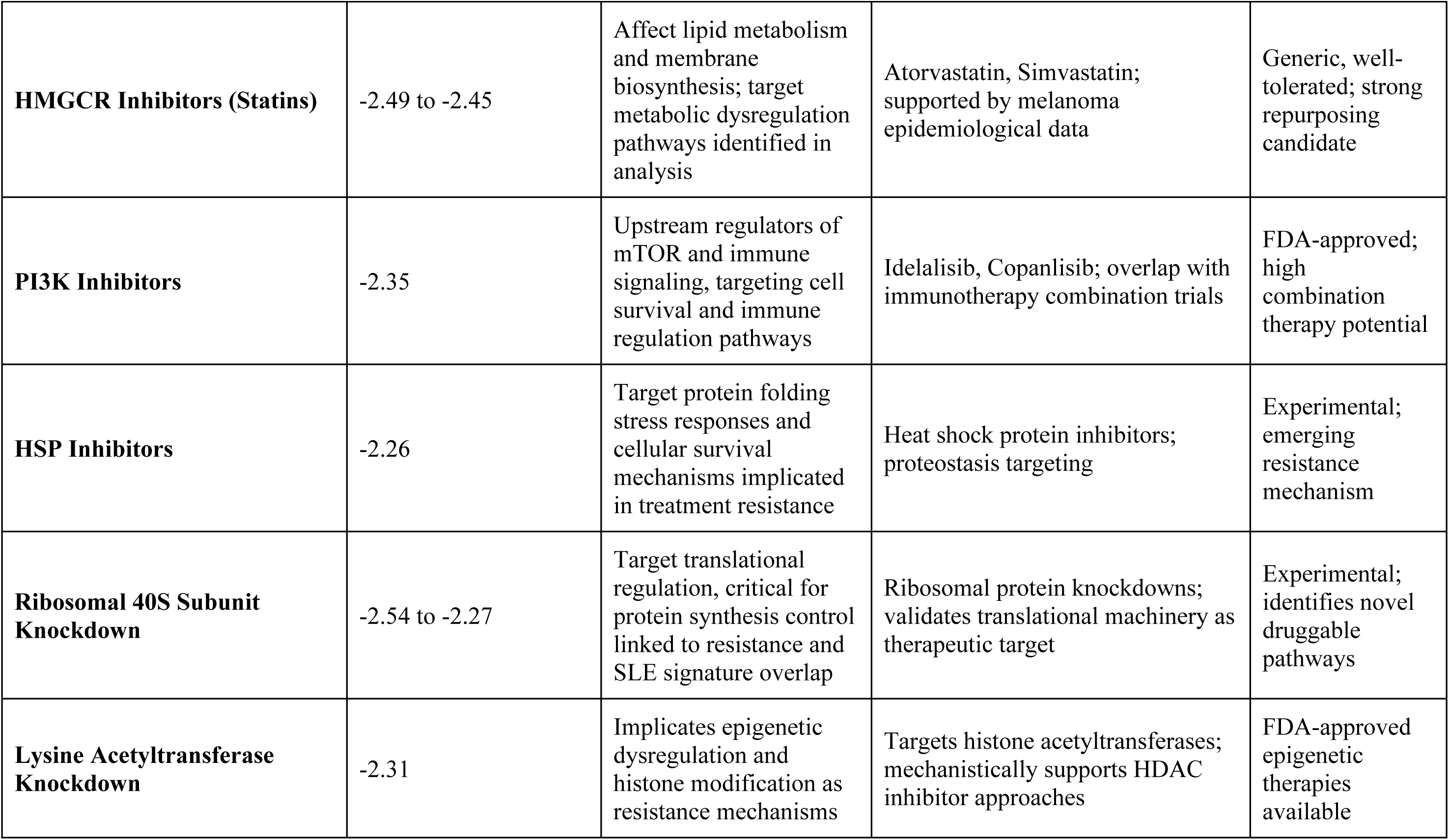
Drug Repurposing Identifies Targeted Candidates

| **Drug / Perturbation Class** | **Connectivity Score Range** | **Mechanistic Fit to Findings** | **Examples / Notes** | **Translational Potential** |
| --- | --- | --- | --- | --- |
| **mTOR Inhibitors** | -2.60 to -2.30 | Target PI3K/mTOR pathway regulating cell cycle progression, protein synthesis, and metabolic dysregulation identified in signatures | Everolimus, Temsirolimus; top scores across multiple cell lines (PC3, HT29) | FDA-approved; active in melanoma immunotherapy combination trials |
| **Proteasome Inhibitors** | -2.58 to -2.29 | Disrupt protein degradation of cyclins/CDKs, modulating cell cycle and immune pathways enriched in non-responders | Bortezomib; supported by preclinical melanoma resistance models and pathway concordance | FDA-approved; experimental combination strategies |
| **Topoisomerase Inhibitors** | -2.50 to -2.28 | Cause DNA damage affecting proliferative cell cycle and DNA repair signatures associated with resistant melanoma | Topotecan, Irinotecan; validated in melanoma cell lines (A375, MCF7) | FDA-approved; applicable in resistant tumor contexts |
| **HMGCR Inhibitors (Statins)** | -2.49 to -2.45 | Affect lipid metabolism and membrane biosynthesis; target metabolic dysregulation pathways identified in analysis | Atorvastatin, Simvastatin; supported by melanoma epidemiological data | Generic, well-tolerated; strong repurposing candidate |
| **PI3K Inhibitors** | -2.35 | Upstream regulators of mTOR and immune signaling, targeting cell survival and immune regulation pathways | Idelalisib, Copanlisib; overlap with immunotherapy combination trials | FDA-approved; high combination therapy potential |
| **HSP Inhibitors** | -2.26 | Target protein folding stress responses and cellular survival mechanisms implicated in treatment resistance | Heat shock protein inhibitors; proteostasis targeting | Experimental; emerging resistance mechanism |
| **Ribosomal 40S Subunit Knockdown** | -2.54 to -2.27 | Target translational regulation, critical for protein synthesis control linked to resistance and SLE signature overlap | Ribosomal protein knockdowns; validates translational machinery as therapeutic target | Experimental; identifies novel druggable pathways |
| **Lysine Acetyltransferase Knockdown** | -2.31 | Implicates epigenetic dysregulation and histone modification as resistance mechanisms | Targets histone acetyltransferases; mechanistically supports HDAC inhibitor approaches | FDA-approved epigenetic therapies available |

Additional promising candidates included topoisomerase inhibitors (−2.50 to -2.28), PI3K inhibitors (−2.35), and HSP inhibitors (−2.26), suggesting multi-pathway targeting approaches. Pathway-level analyses revealed that ribosomal subunit knockdown (−2.54), proteasome pathway disruption (−2.30), and cell cycle inhibition (−2.30) strongly reversed the signature, supporting the mechanistic relevance of these therapeutic targets. These findings require preclinical validation to confirm efficacy and optimal combination strategies.

## Conclusion

This study identifies convergent yet disease-specific molecular pathways linking systemic lupus erythematosus (SLE) and melanoma immunotherapy response. The immune signature (R_up ∩ SLE), enriched in interferon signaling and epigenetic regulators such as ASF1B and EZH2, delineates a shared framework whereby chromatin remodeling facilitates robust immune activation in both autoimmunity and checkpoint blockade response[4,7,11,58]. In contrast, the cell cycle signature (NR_up ∩ SLE), characterized by proliferation drivers including CDK1 and CCNB1, emerges as a hallmark of immune checkpoint blockade resistance and poor prognosis in melanoma, with limited relevance in systemic autoimmunity [39,53] findings highlight that immune activation constitutes a convergent biological feature, whereas proliferative programs are largely disease-specific.

The cell cycle signature demonstrated strong prognostic value in melanoma survival analyses, underscoring its potential utility as a biomarker for adverse outcomes[17,40]. Meanwhile, the immune signature showed promising, though statistically non-significant, protective trends, warranting further validation in immune checkpoint blockade-treated cohorts. Connectivity Map analysis nominated therapeutics that target these epigenetic and proliferative pathways—including mTOR inhibitors, proteasome inhibitors, HDAC inhibitors, and statins[43,50]—with statins representing a novel, cost-effective candidate supported by epidemiological data [47–50]. Nonetheless, experimental and clinical validation remain essential to confirm these candidates’ efficacy and optimal therapeutic contexts.

Overall, our integrative analysis elucidates epigenetic remodeling as a critical mechanistic bridge connecting autoimmune dysregulation and melanoma immunotherapy response, offering actionable biomarker candidates and rational therapeutic avenues for future translational and preclinical investigation.

### Limitations

This study provides novel insights into convergent pathways between systemic lupus erythematosus (SLE) and melanoma, yet several limitations should be acknowledged. First, the immune signature’s non-significant prognostic association in the TCGA-SKCM cohort (HR = 0.259, p = 0.121) likely reflects the limitations of bulk RNA-sequencing and the predominance of untreated patients in this dataset, which may obscure immune signals relevant to immune checkpoint blockade (ICB) [40]. This restricts the generalizability of the immune signature as a biomarker outside ICB-treated contexts. Second, the moderate discriminatory performance of the signatures (AUC = 0.6567–0.780 across validation datasets) indicates that larger and more homogeneous cohorts are needed to improve biomarker precision [17, 18]. Third, the reliance on transcriptomic analyses without functional validation (e.g., CRISPR knockdown of hub genes such as EZH2 or CDK1) limits mechanistic interpretation of epigenetic and cell cycle pathways [7, 39,61]. Fourth, the limited relevance of the cell cycle signature in SLE (FDR-adjusted p = 0.2414 in GSE261866) reflects disease-specific differences: systemic autoimmunity in SLE contrasts with the tumor-immune microenvironment in melanoma, complicating direct cross-disease comparisons [12, 62]. Finally, the drug repurposing analysis—although supported by preclinical and early clinical evidence [45–50]—remains hypothesis-generating, as no in vitro or in vivo validation was performed for candidates such as mTOR inhibitors, proteasome inhibitors, HDAC inhibitors, or statins.

### Future Directions

Future work should prioritize validating the immune and cell cycle signatures in larg ICB-treated melanoma cohorts to more directly assess immunotherapy-relevant biology [17, 40]. Integrating multi-omics data, including epigenomics and proteomics, may better capture disease-specific mechanisms, particularly the role of epigenetic regulators such as ASF1B and EZH2 in SLE and melanoma [7, 44]. Functional experiments using CRISPR-mediated knockdown or overexpression of hub genes (e.g., CDK1, EZH2) are critical to confirm their roles in immune activation and resistance to ICB [39]. Preclinical testing of therapeutic candidates (mTOR inhibitors, proteasome inhibitors, HDAC inhibitors, and statins) in SLE and melanoma models— especially in combination with ICB—could help establish their capacity to enhance immunotherapy efficacy or modulate autoimmunity [45–50]. Given epidemiological evidence, statins in particular warrant further exploration as low-cost, immunomodulatory agents across both diseases [47, 50]. Finally, larger, well-characterized cohorts with single-cell RNA-sequencing would refine biomarker accuracy and delineate cell-type-specific contributions, bridging the gap between systemic autoimmunity and the tumor-immune microenvironment [6, 12]. These efforts could ultimately translate convergent immune–epigenetic pathways into clinically actionable biomarkers and therapeutic strategies.

---

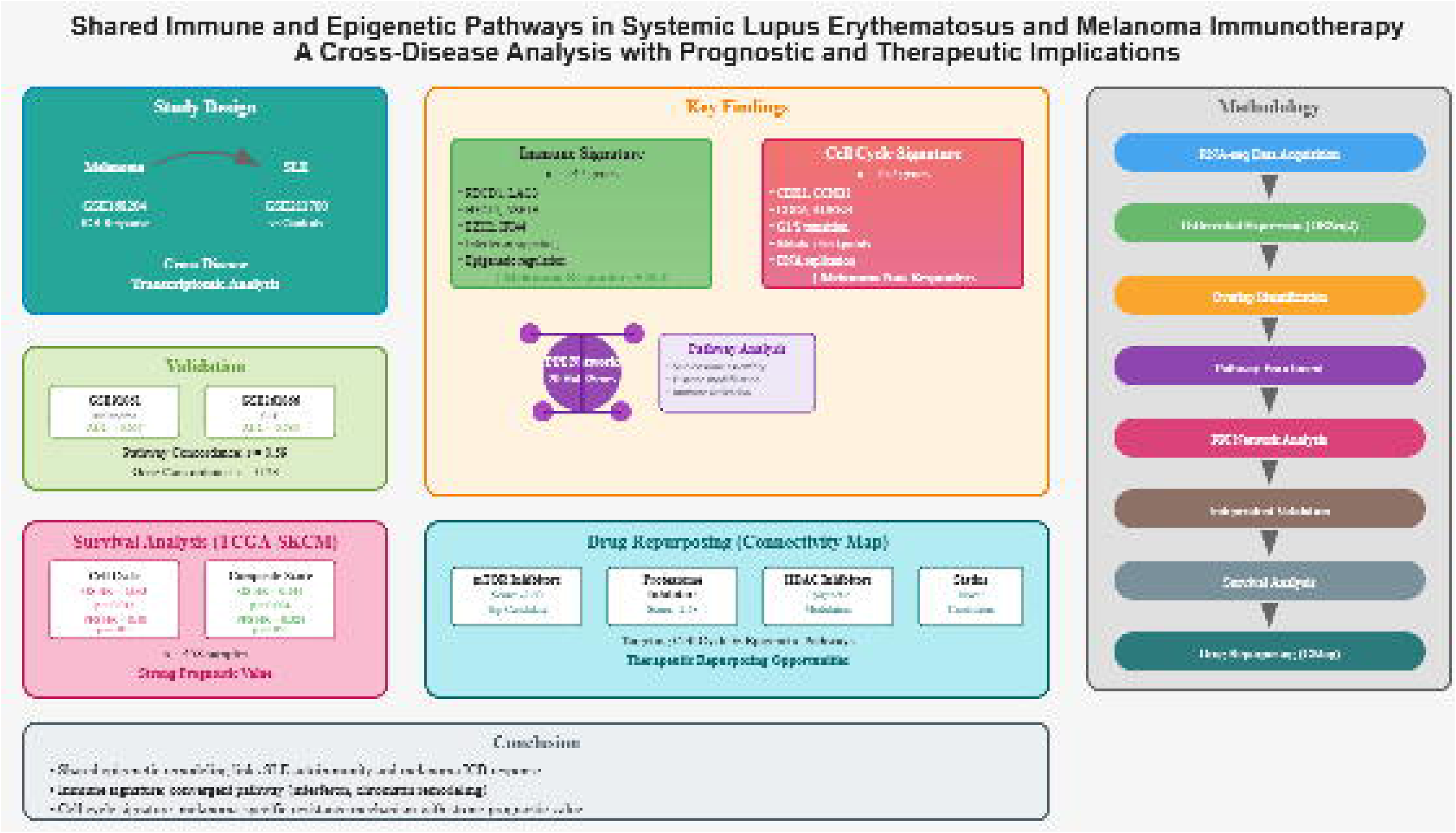

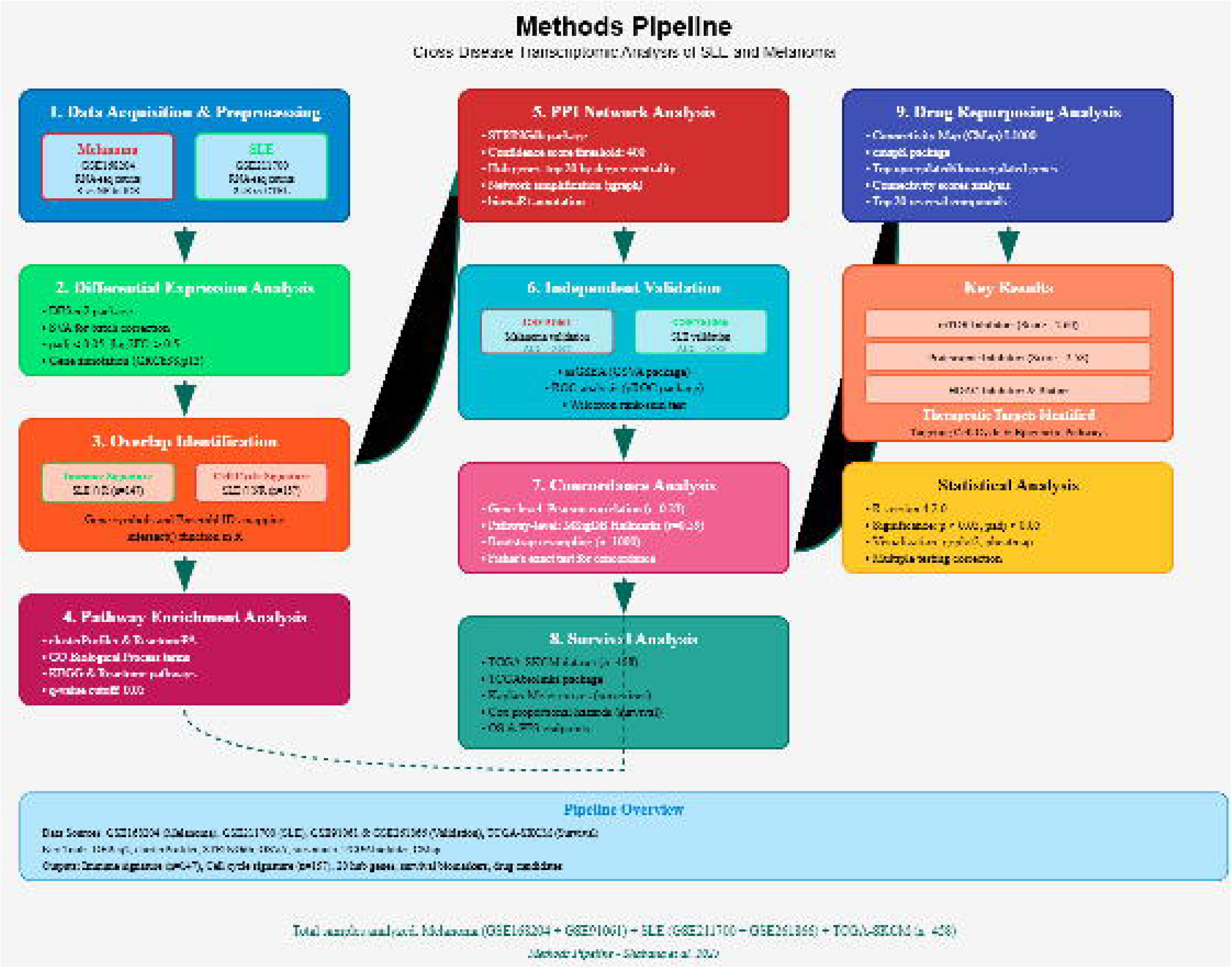

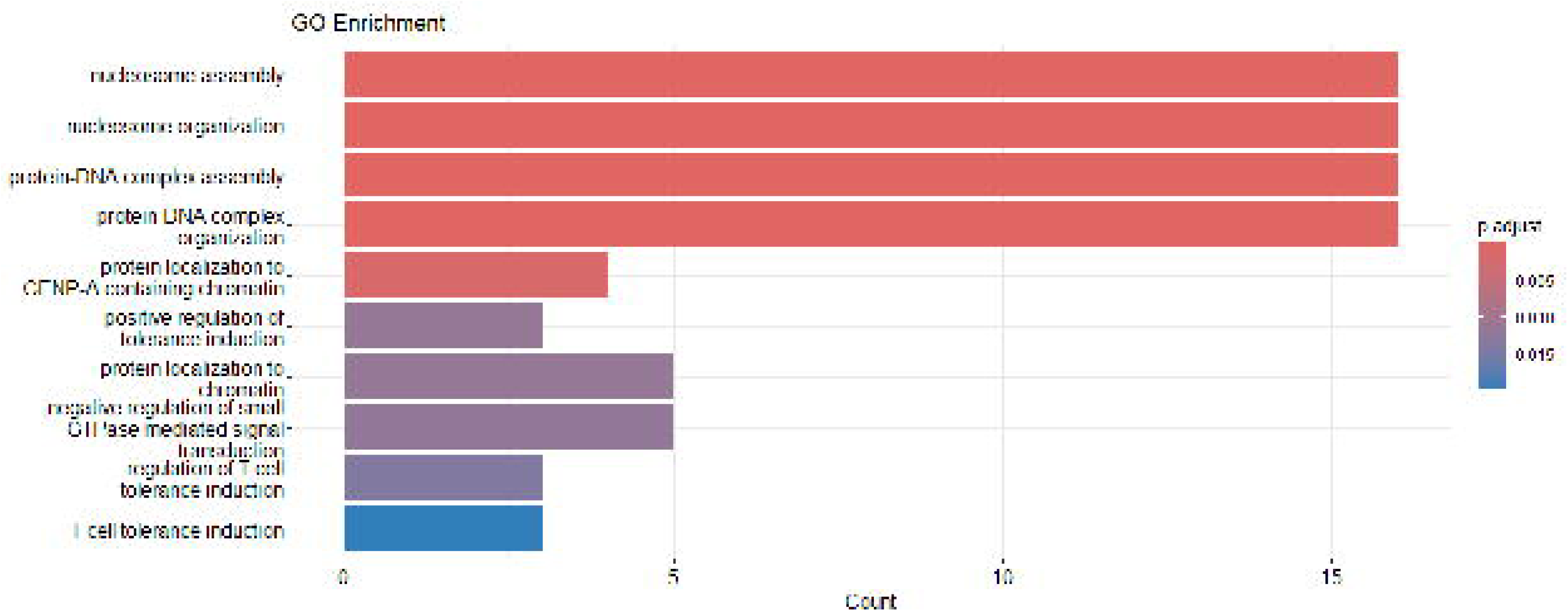

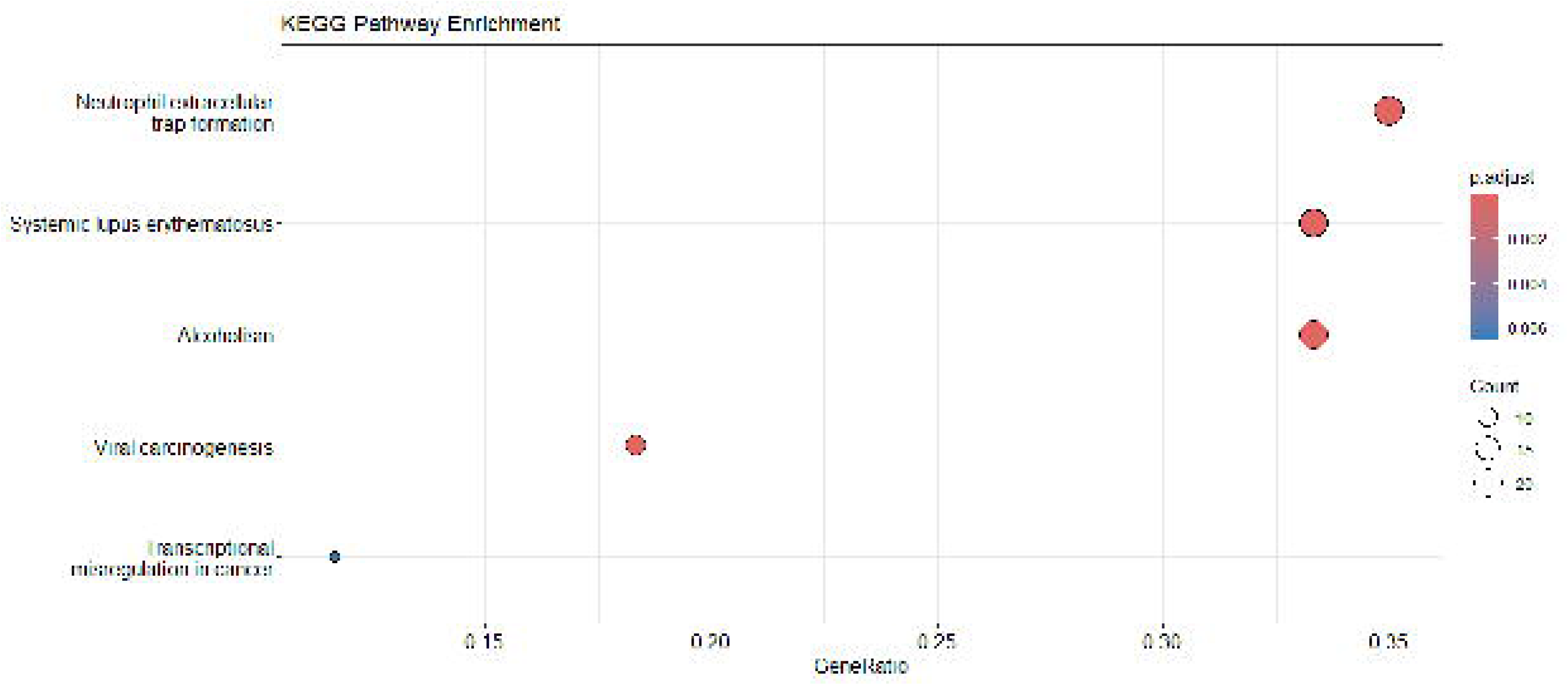

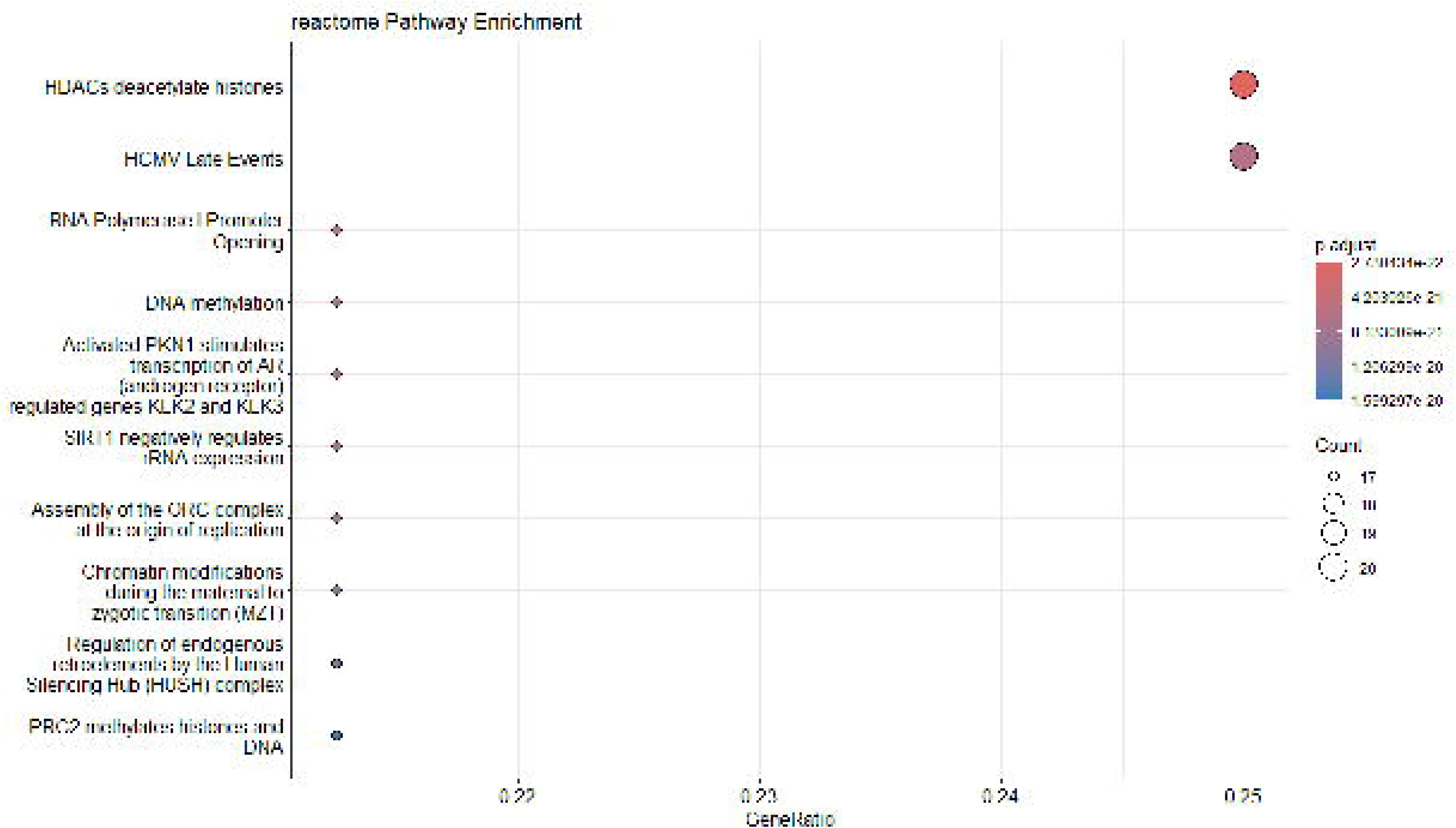

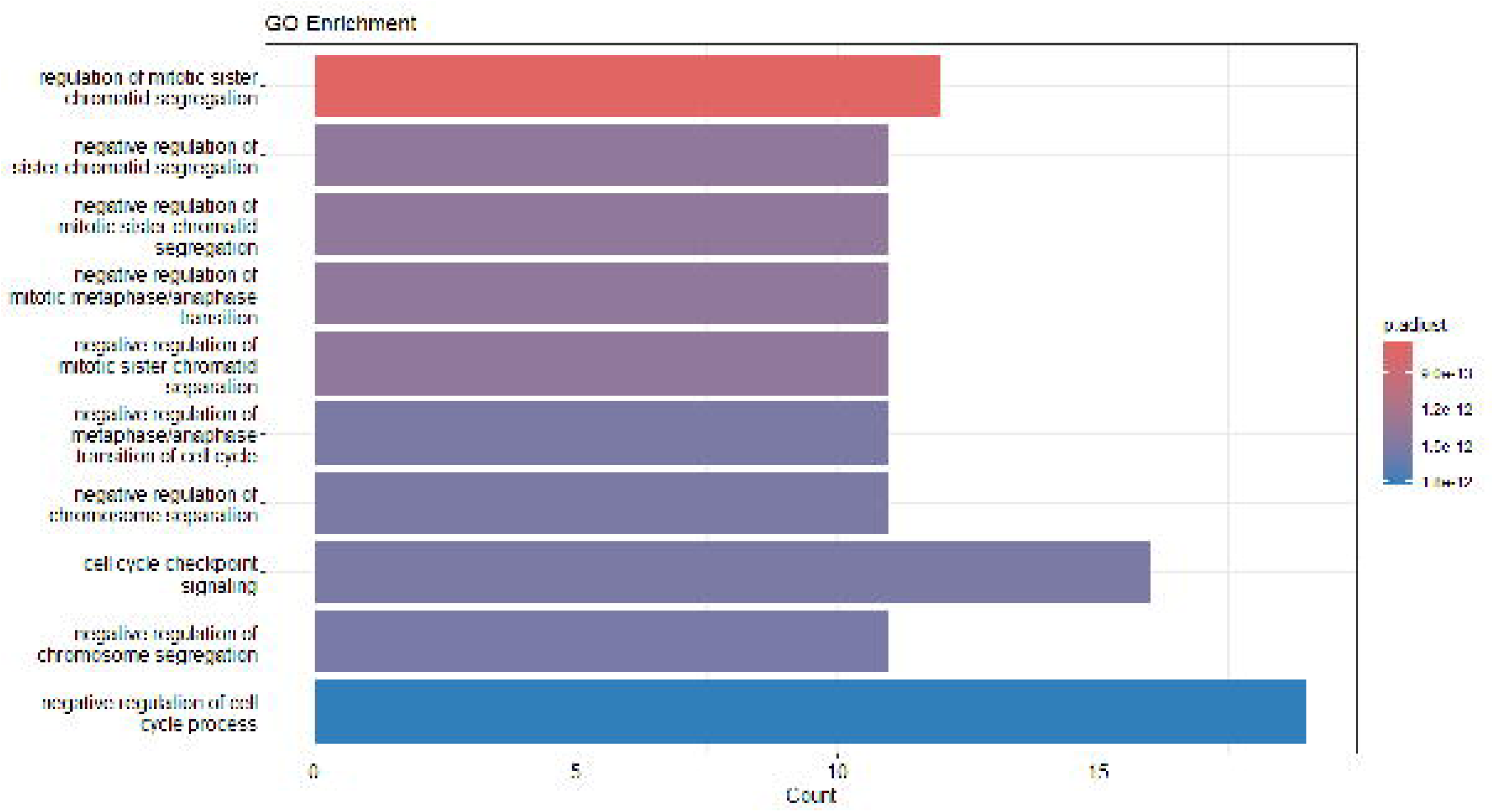

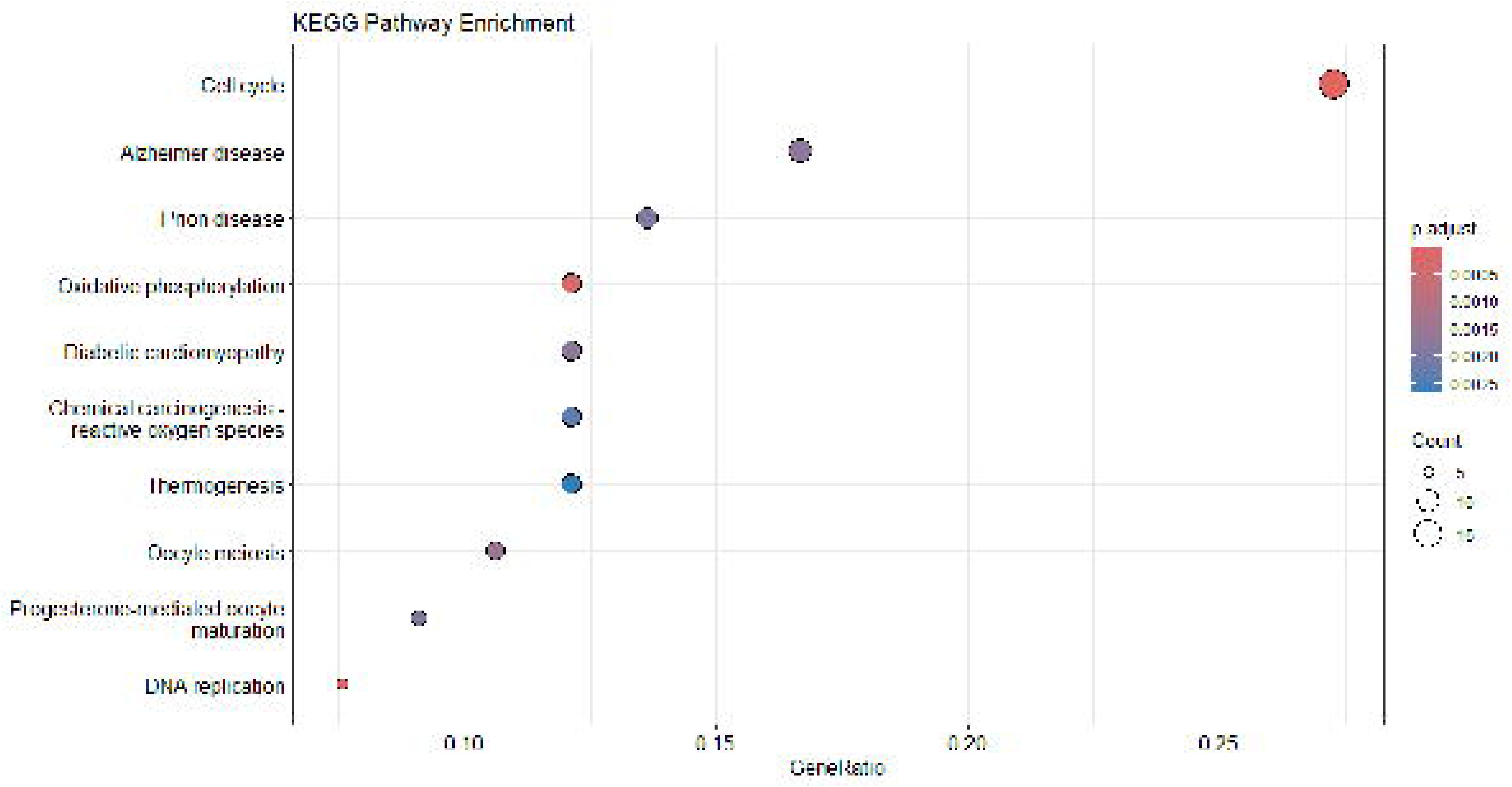

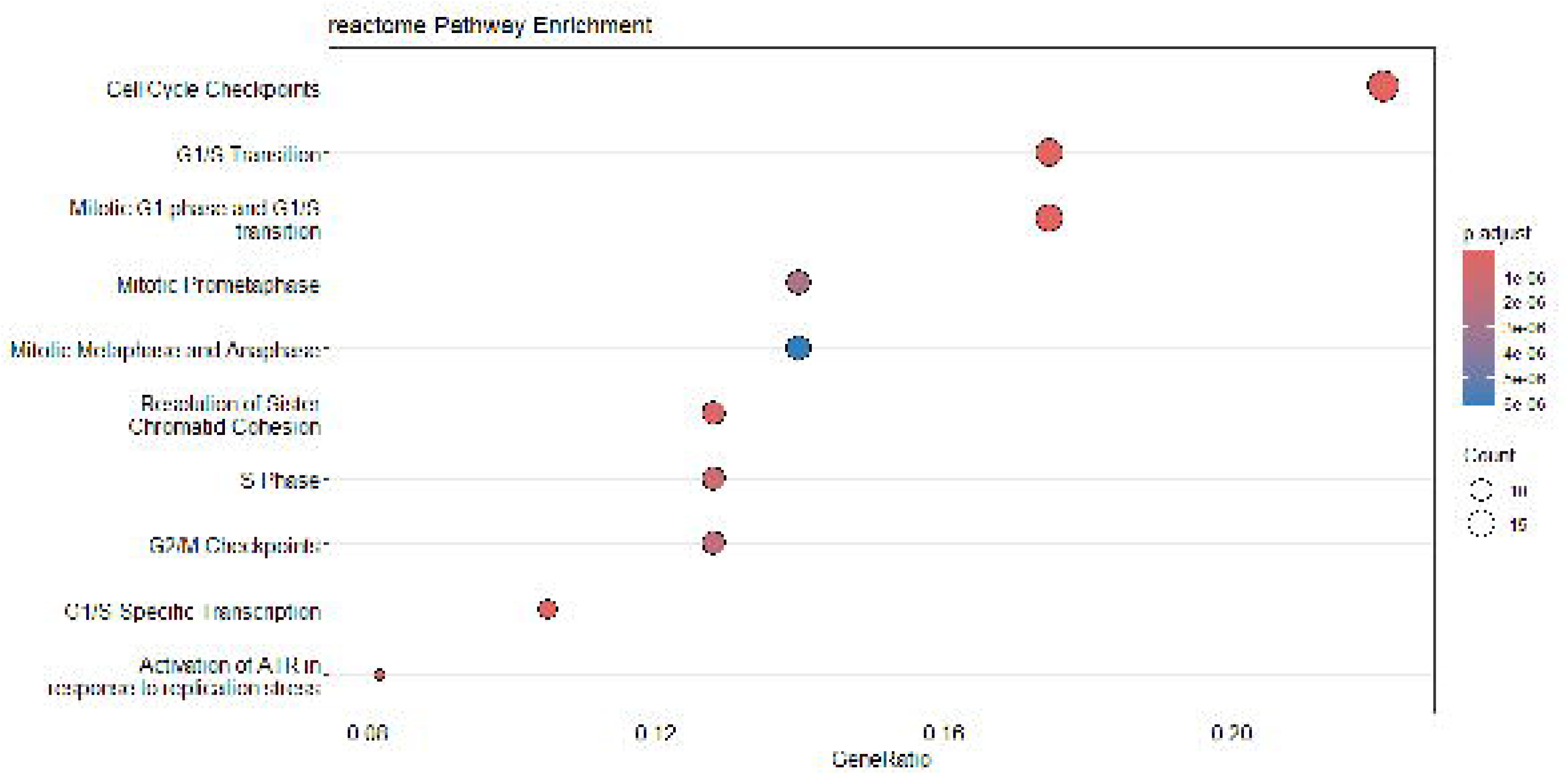

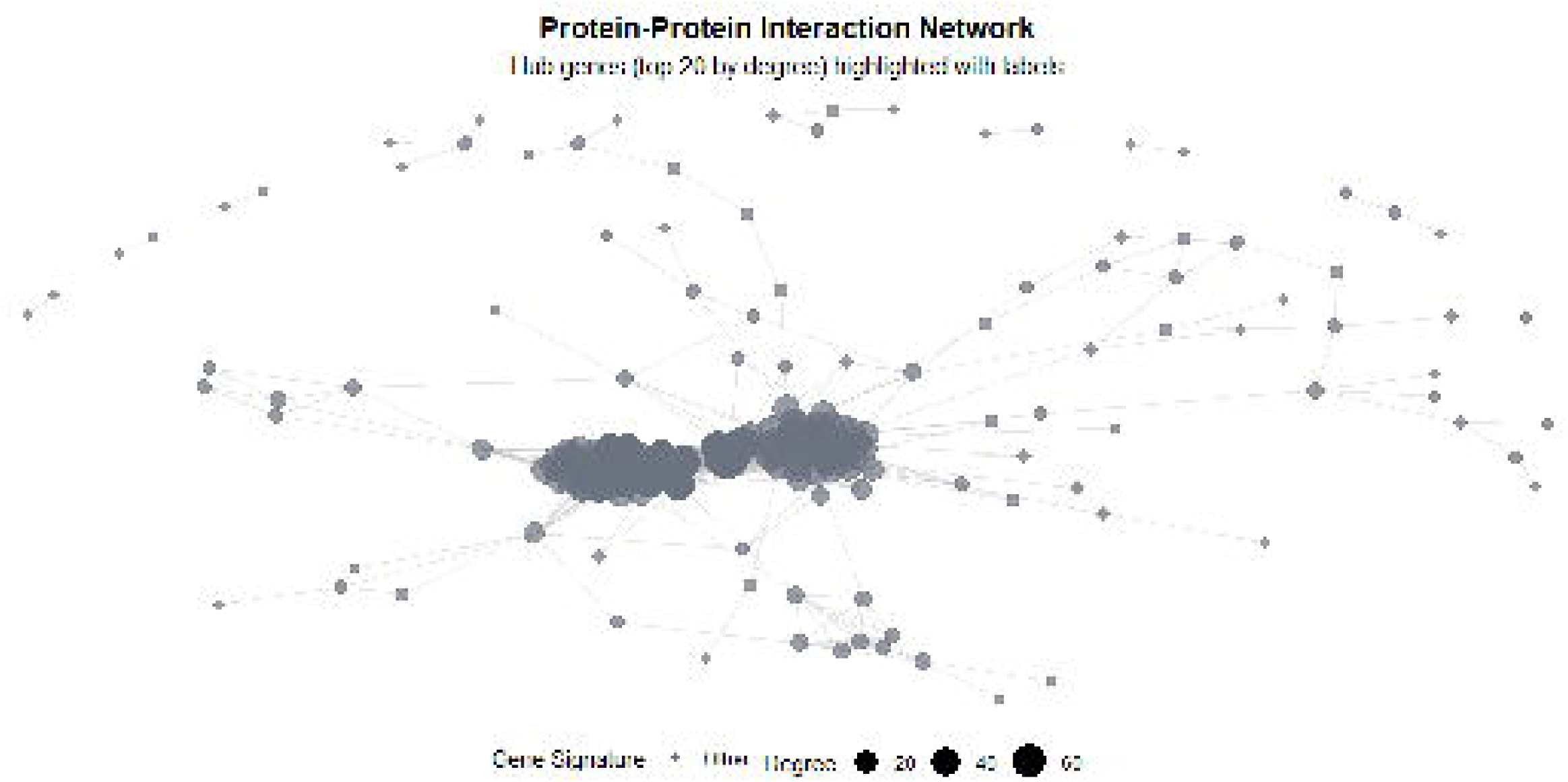

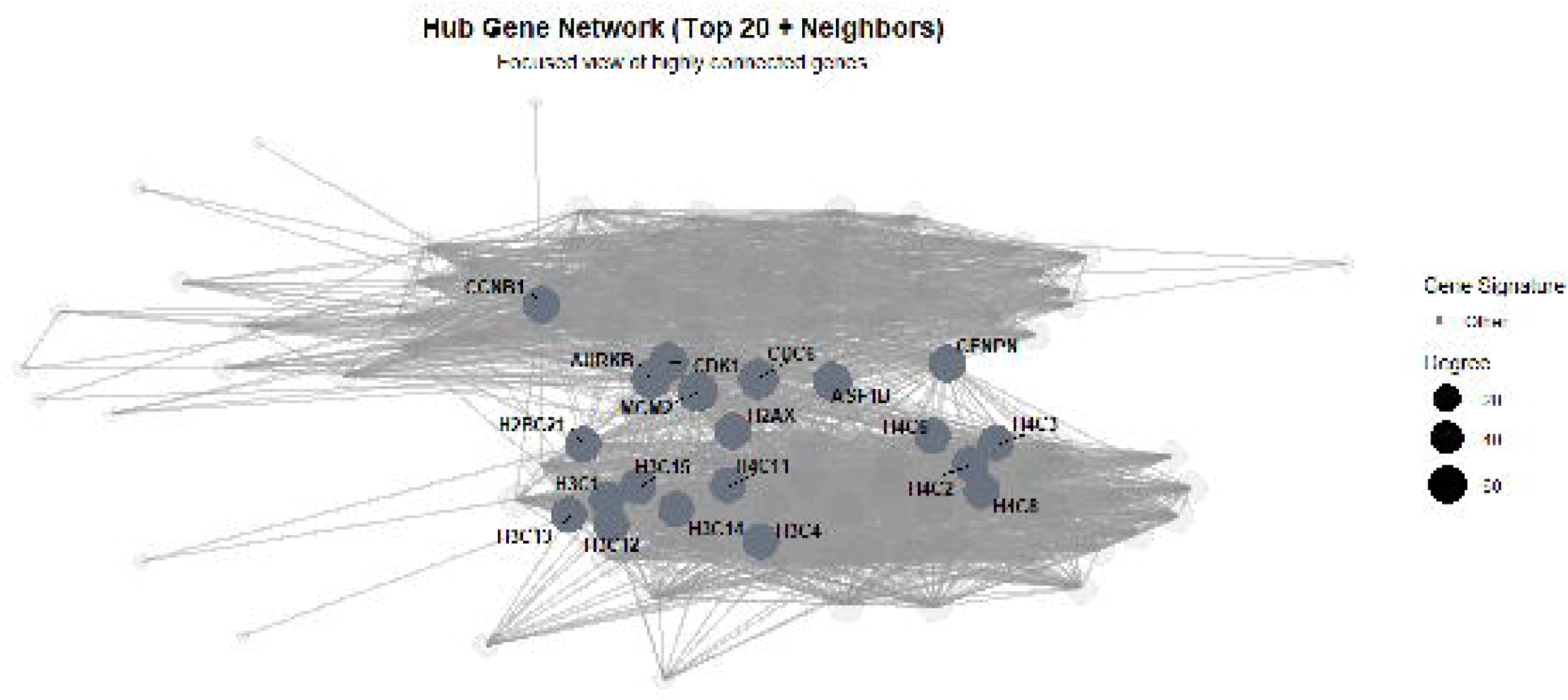

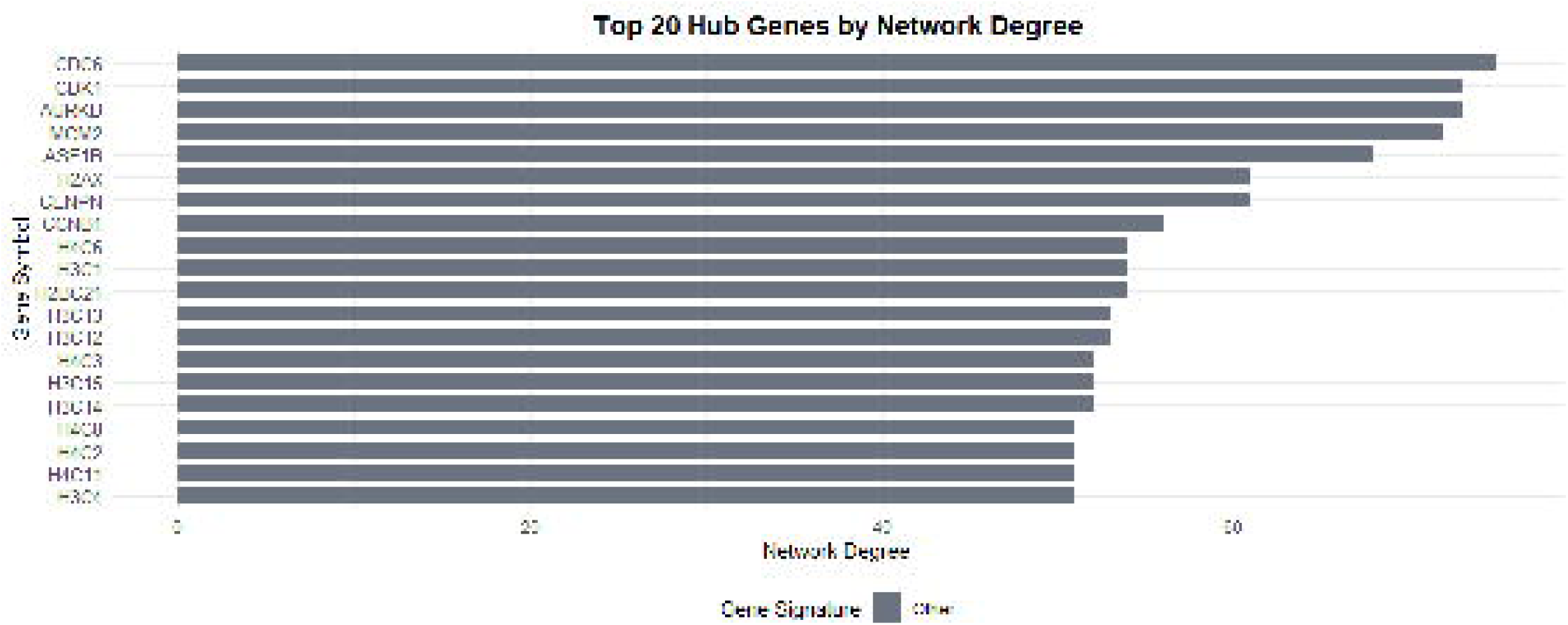

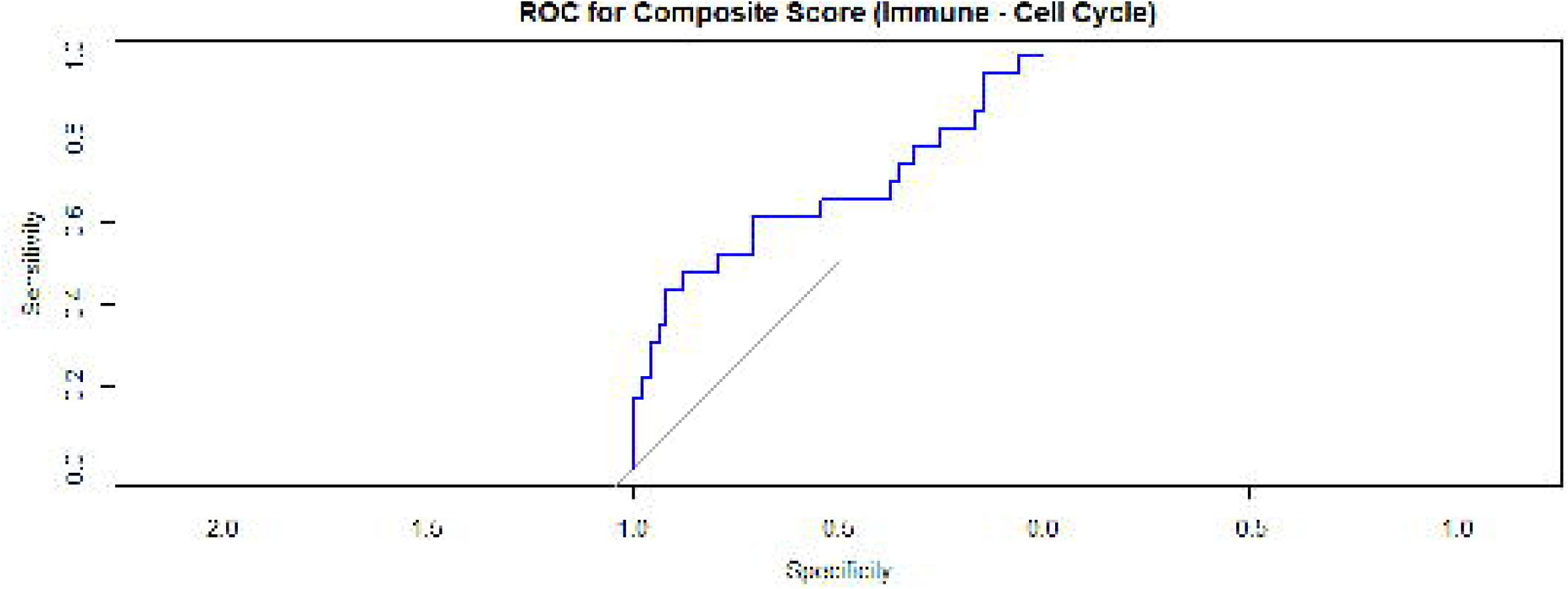

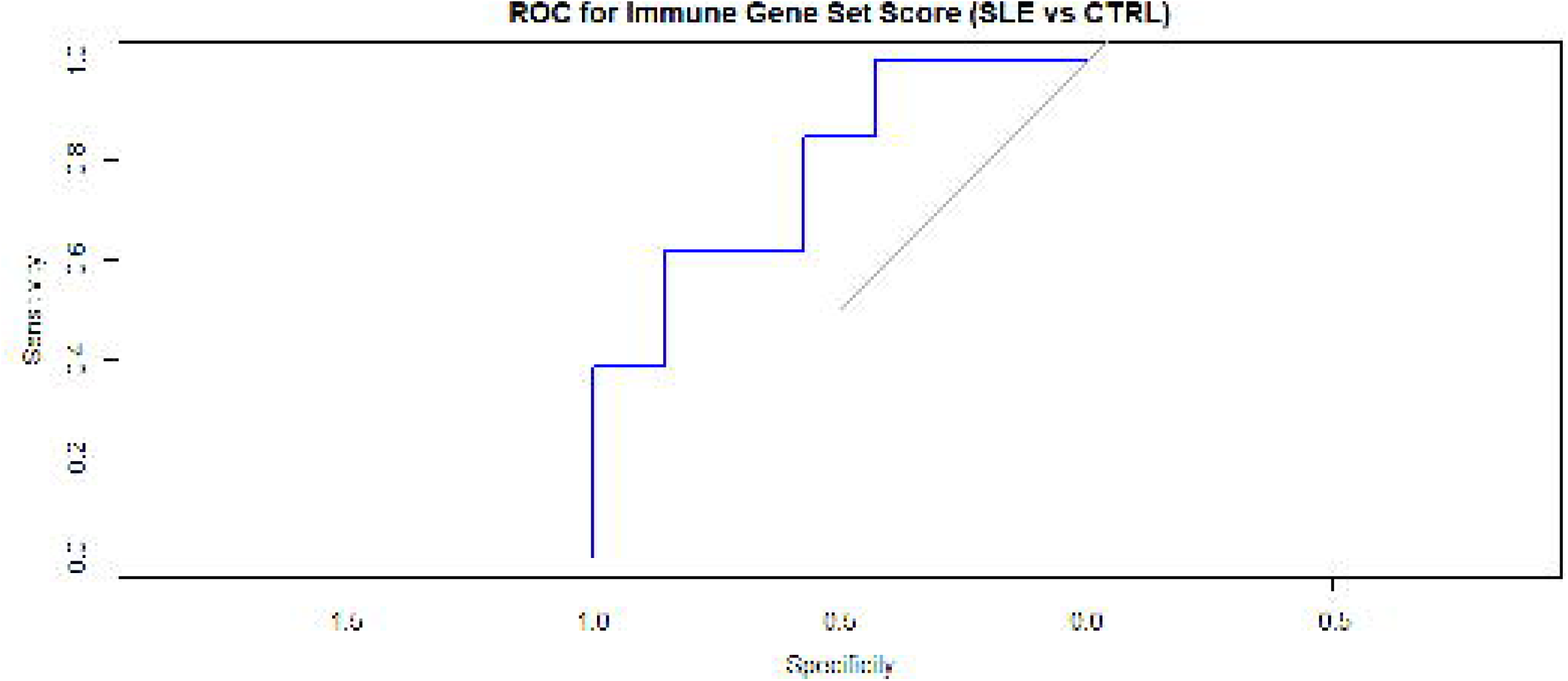

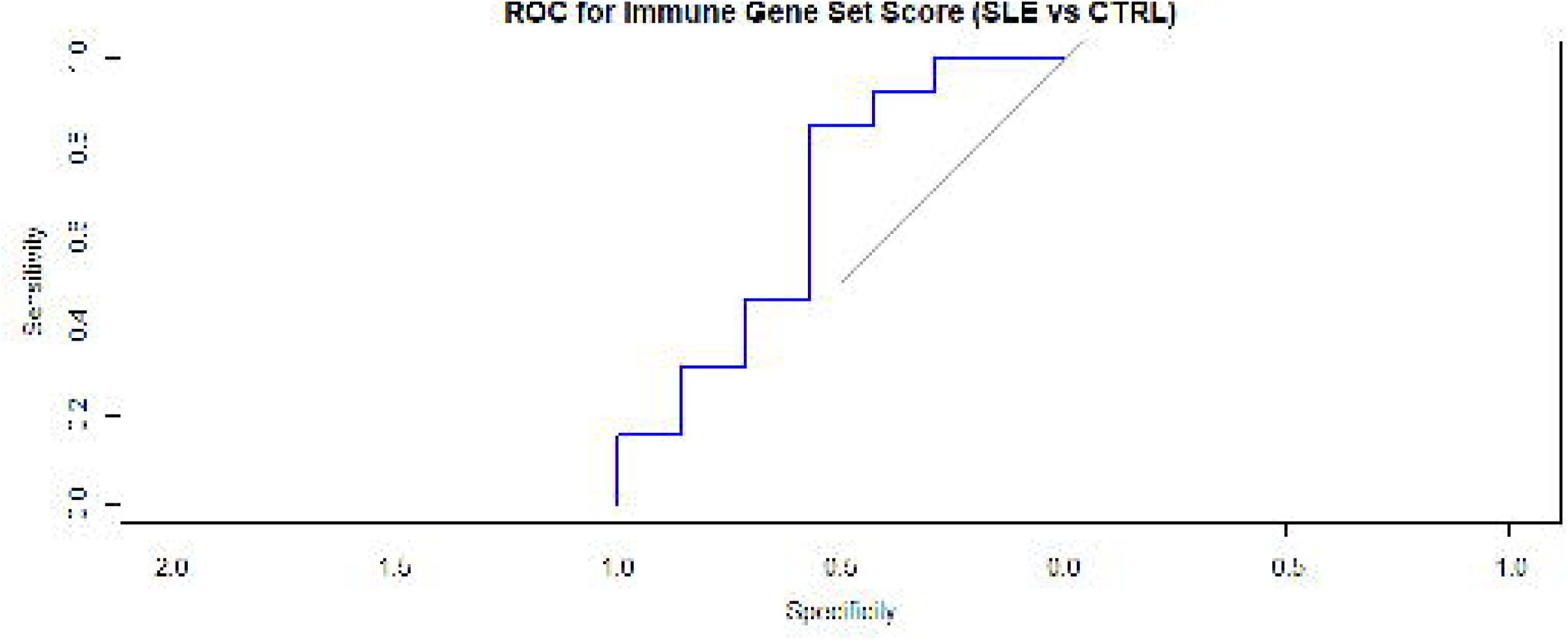

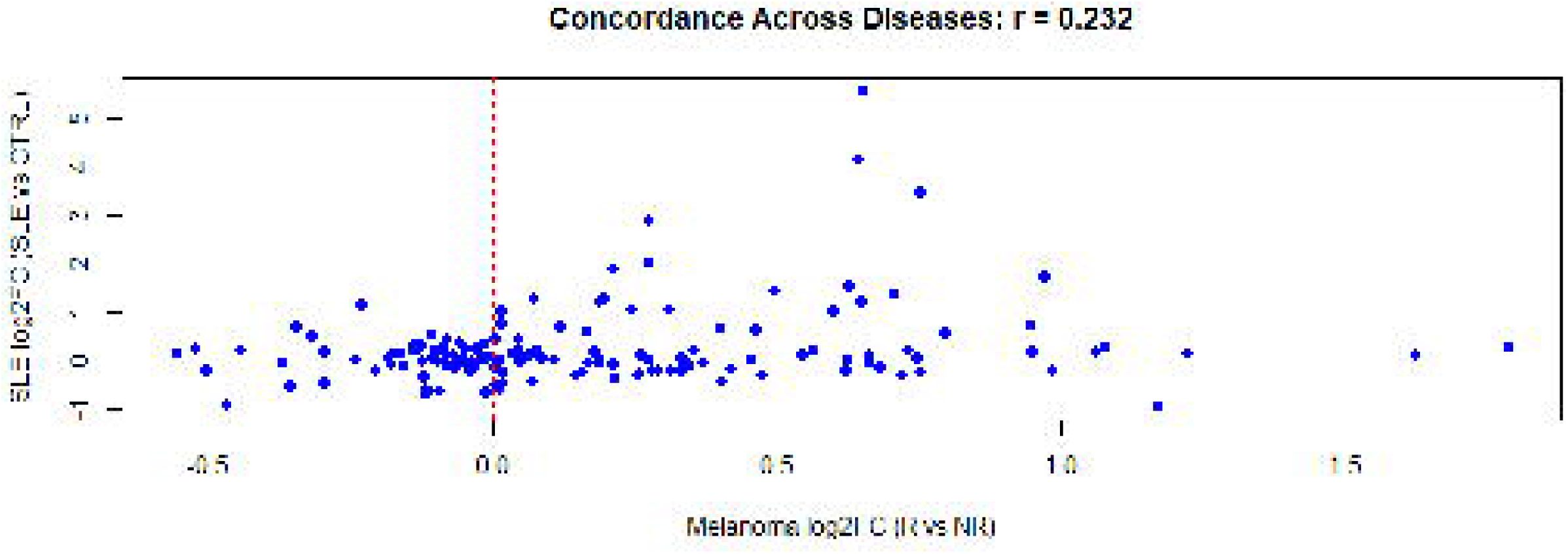

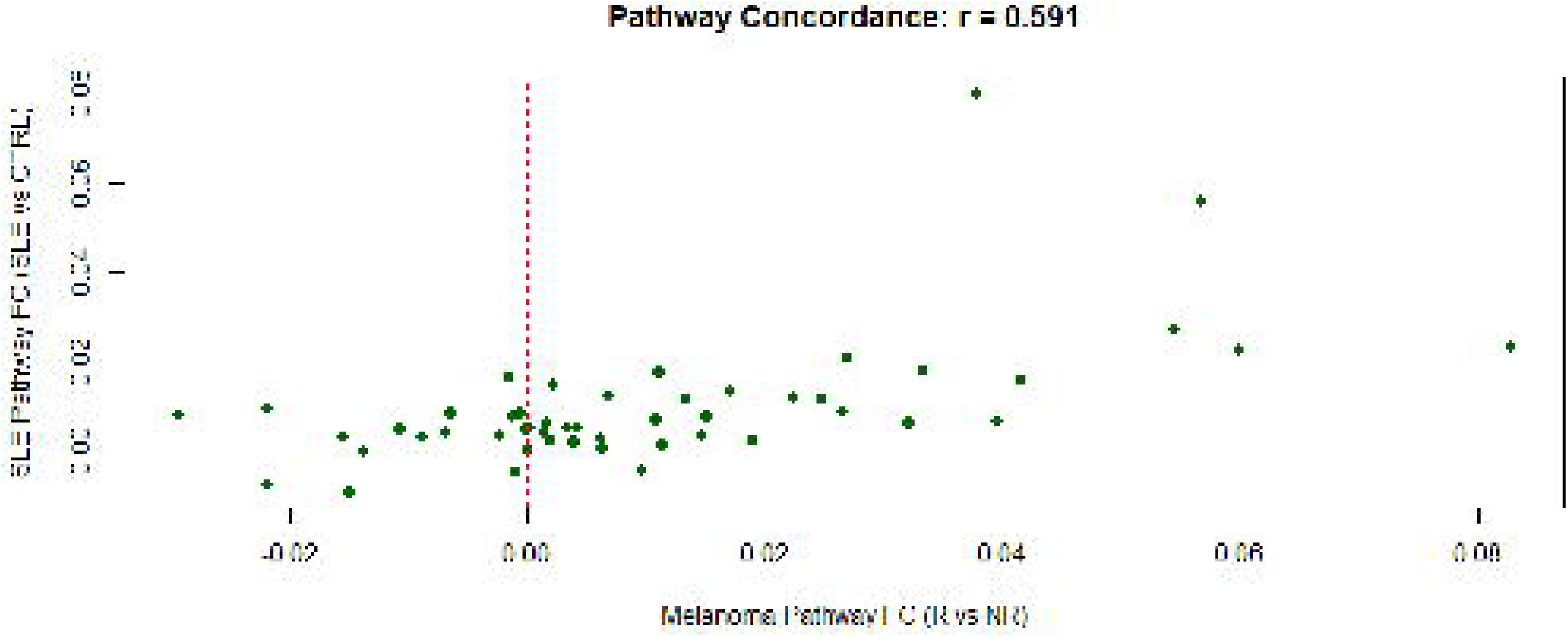

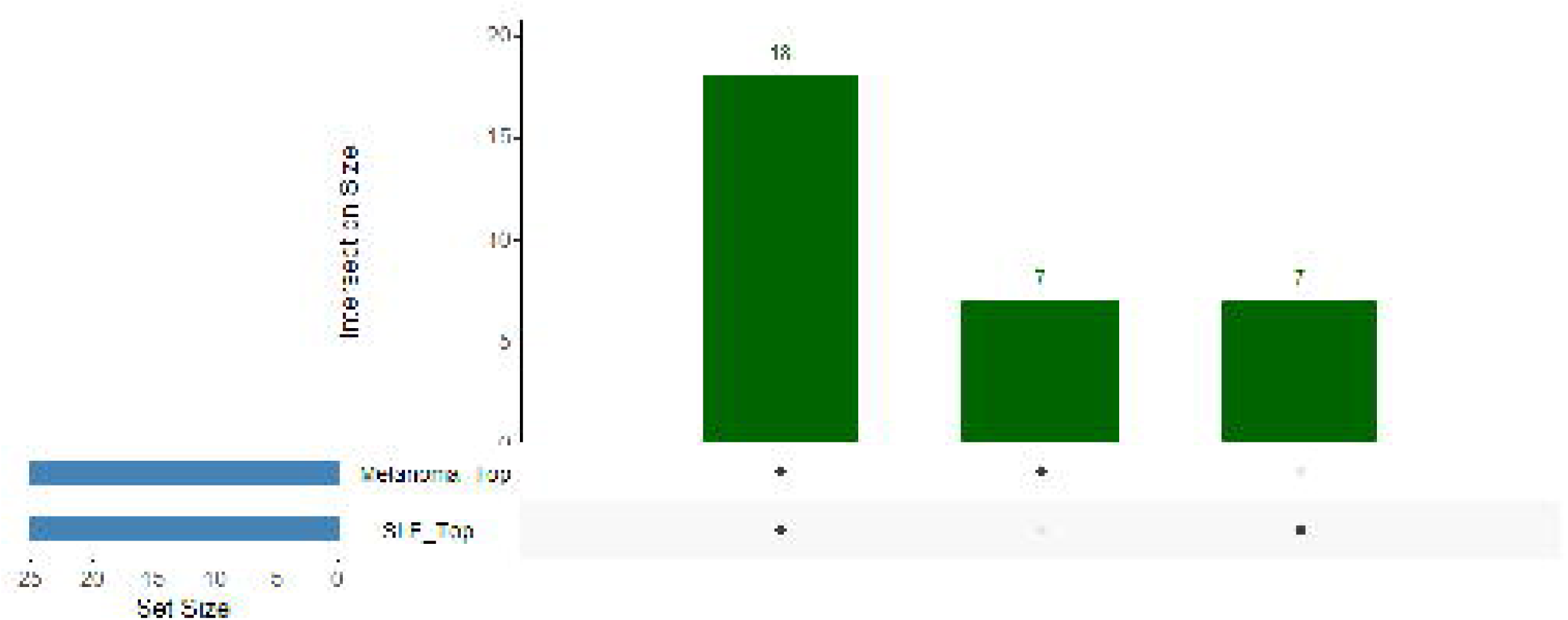

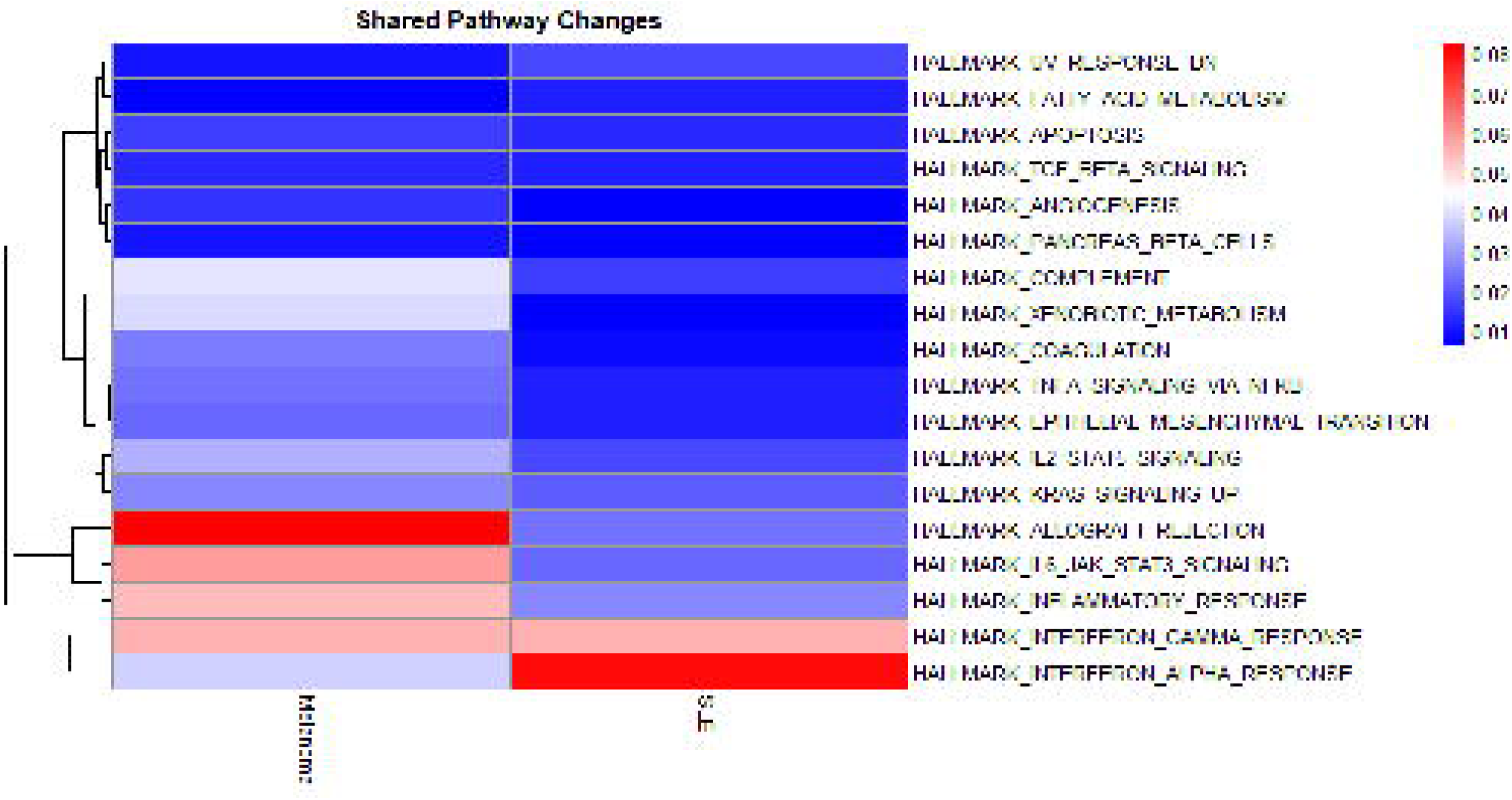

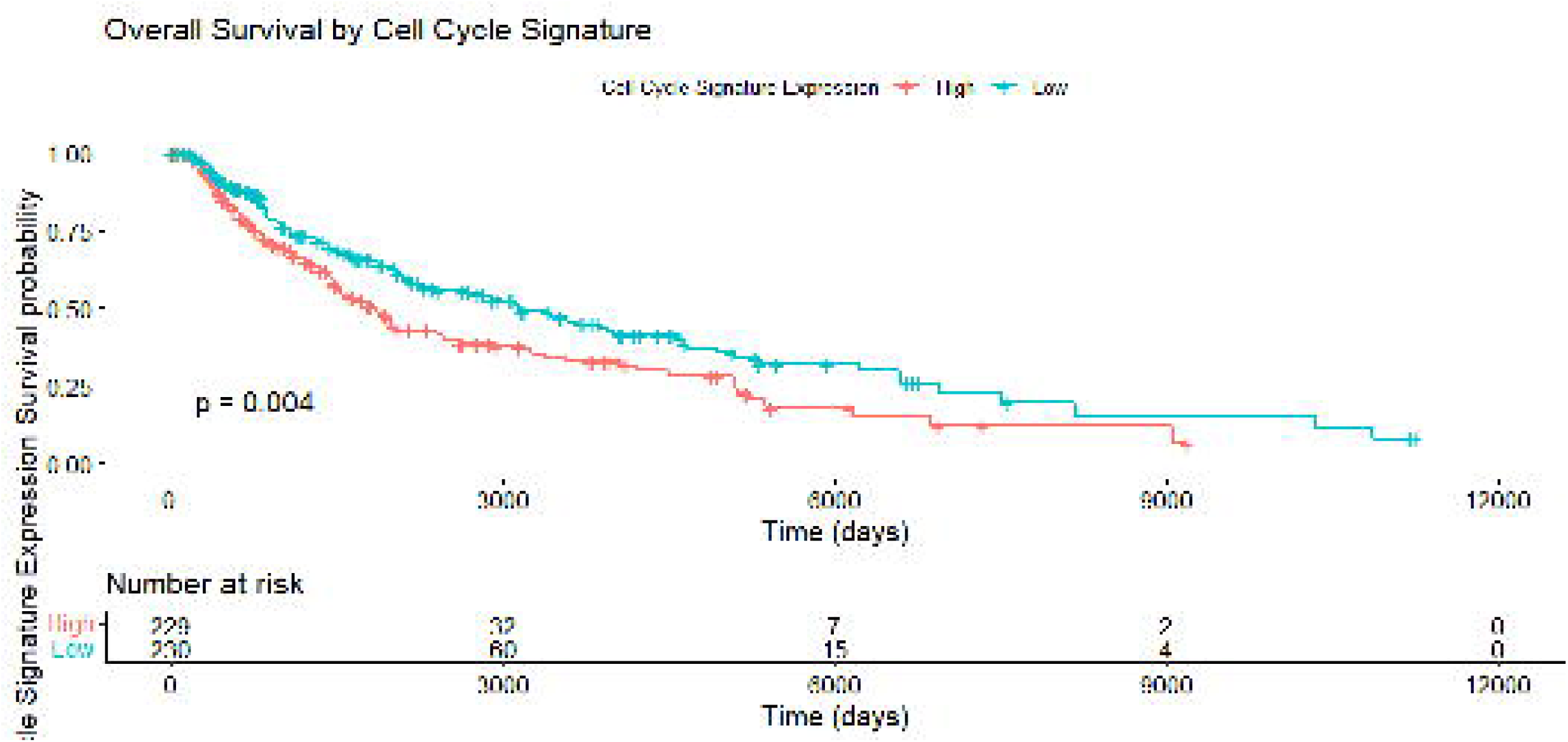

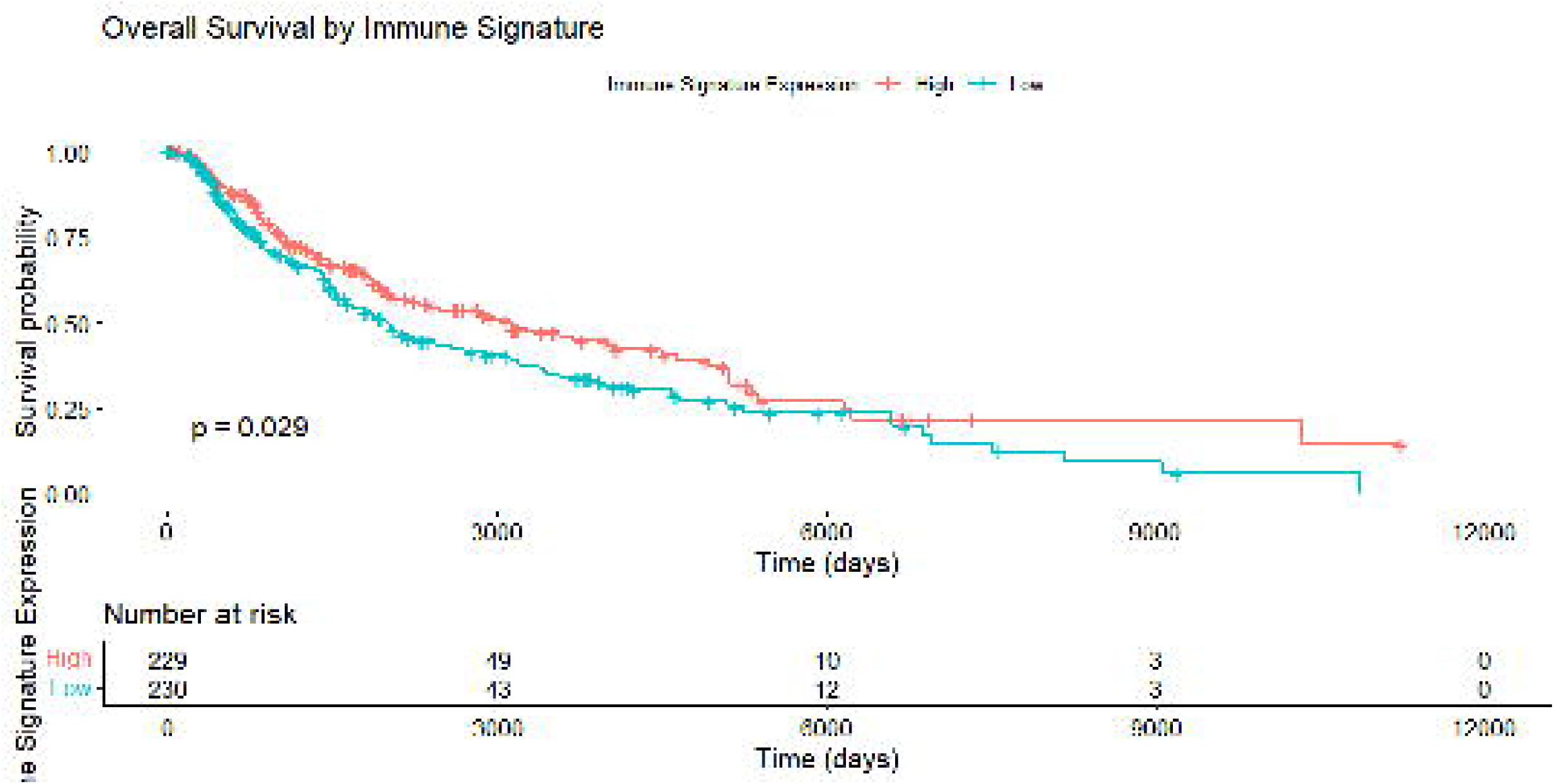

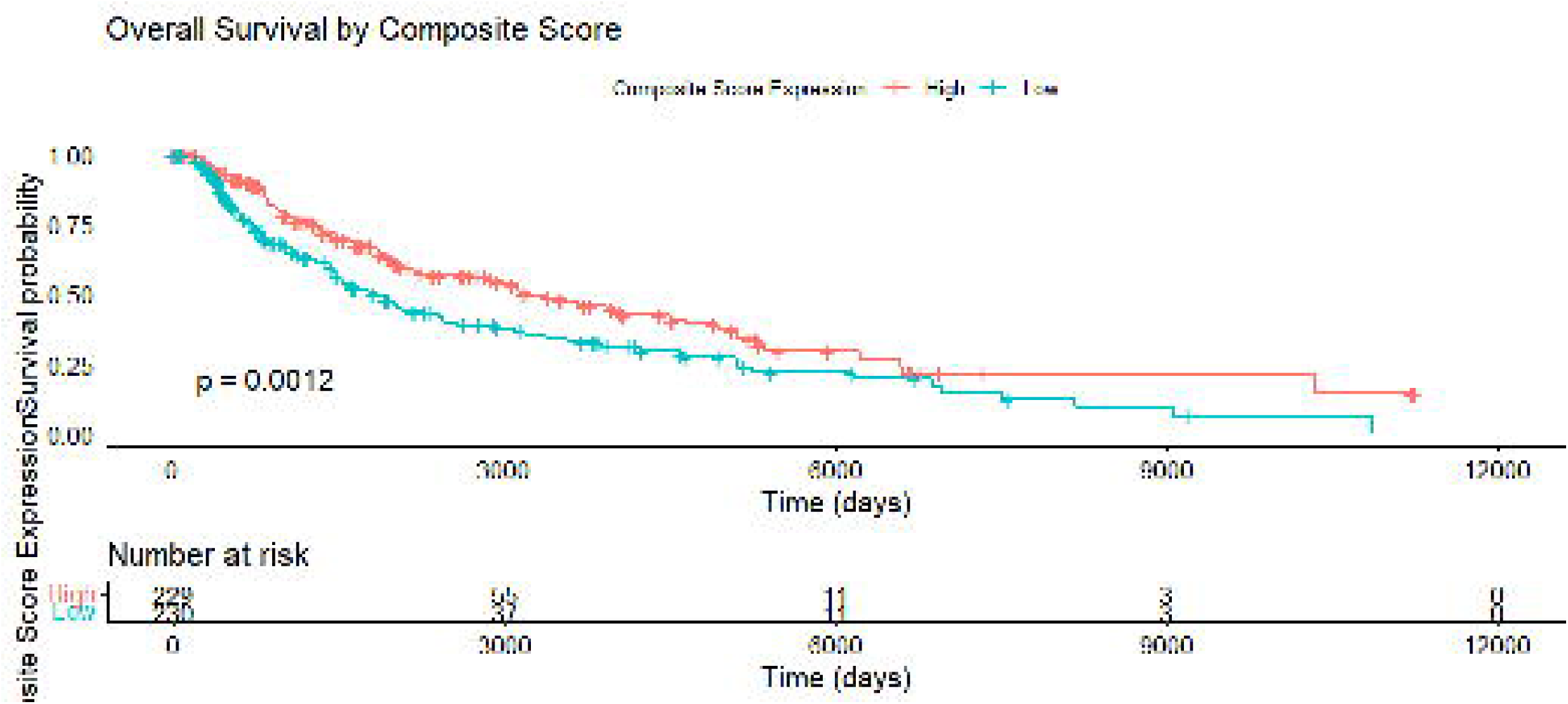

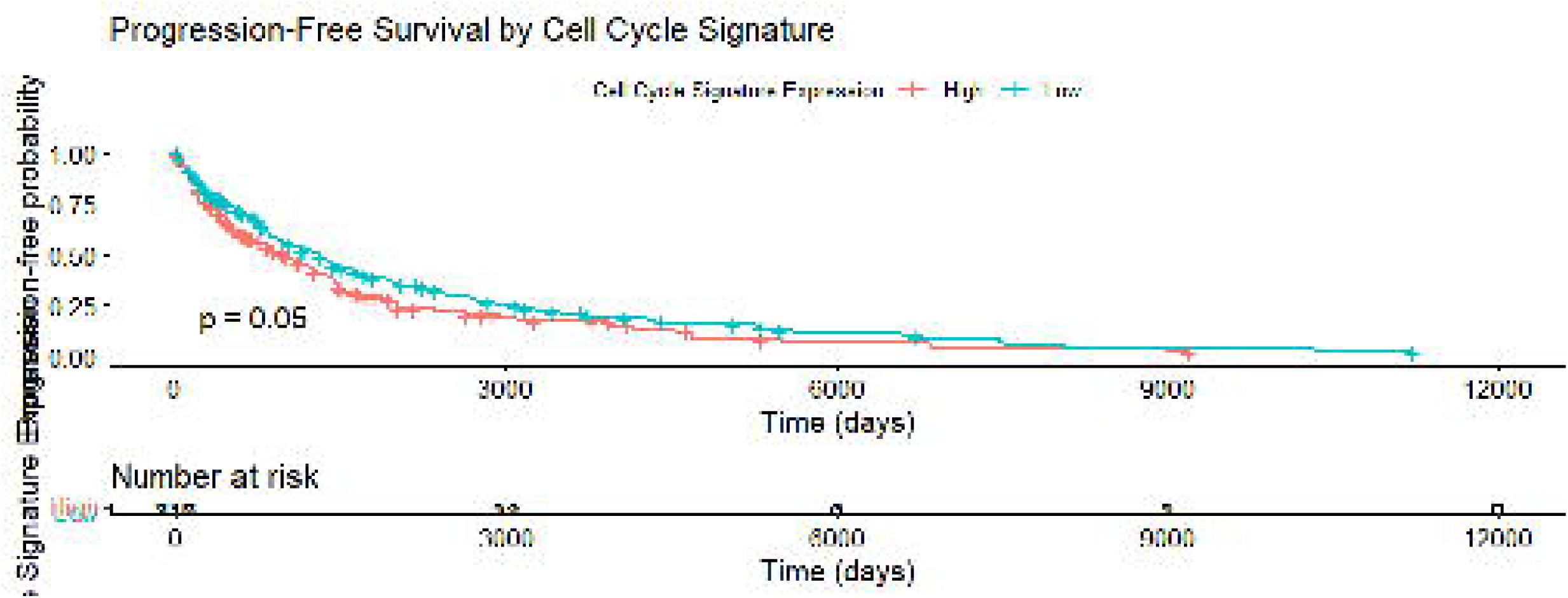

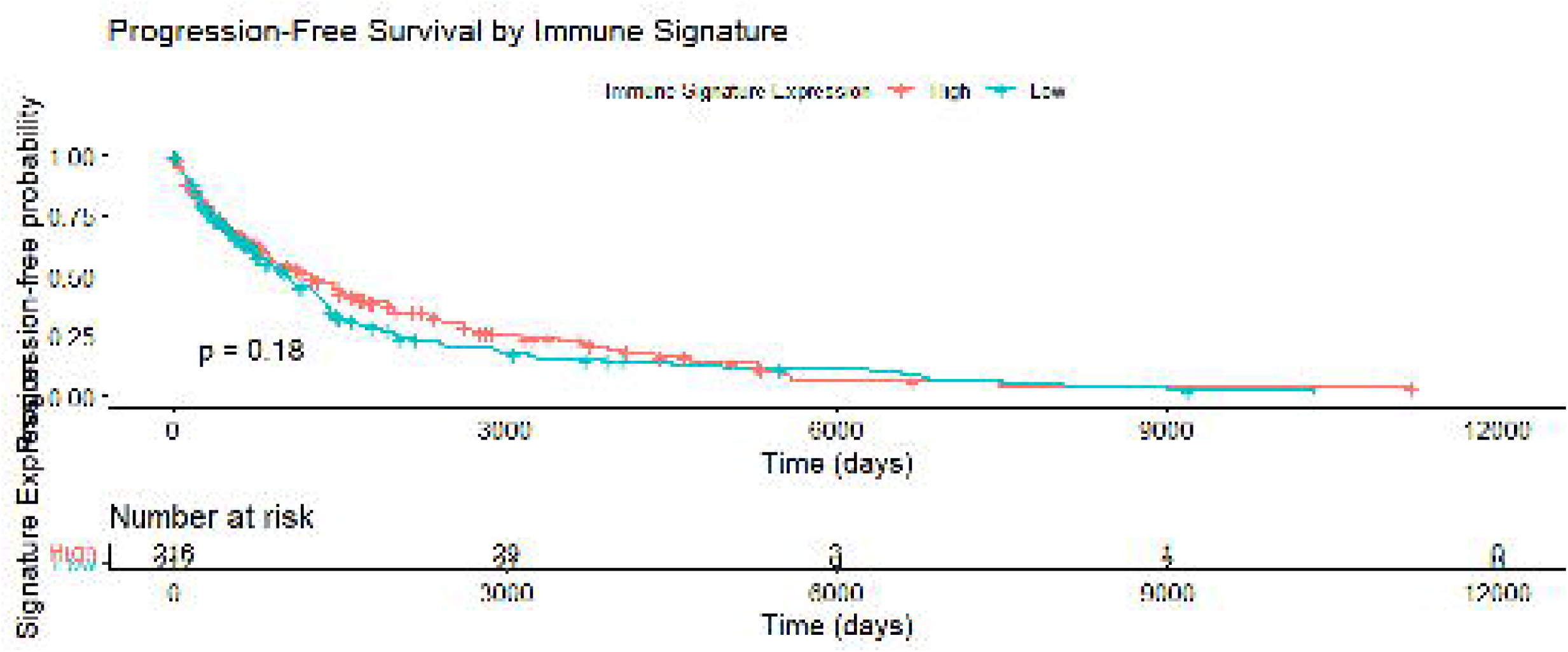

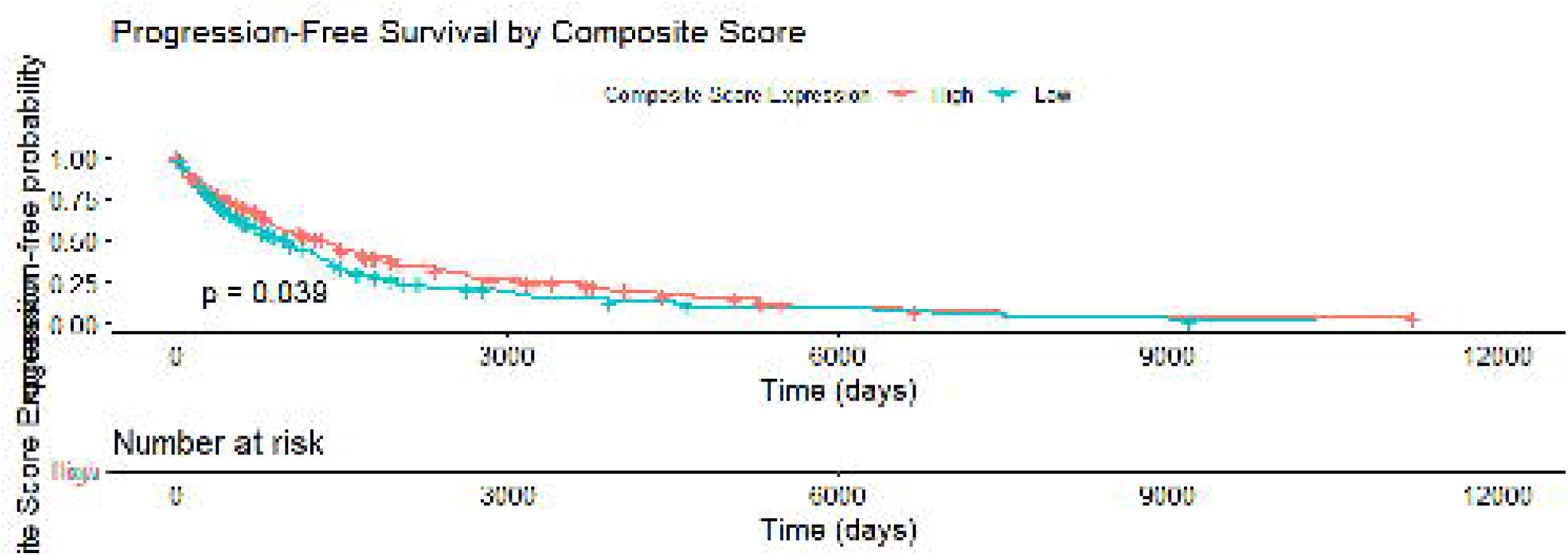

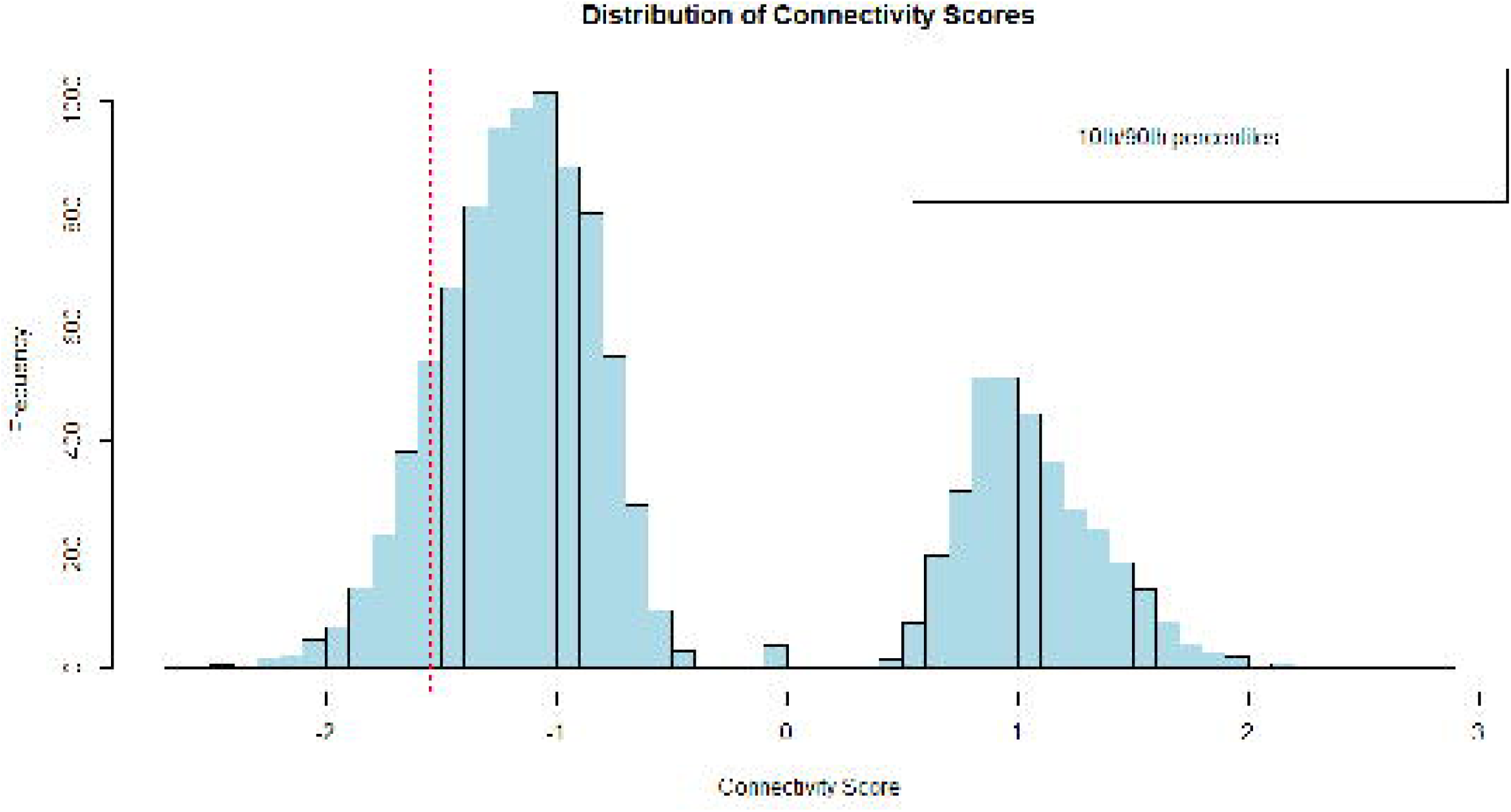

